# Mechanical stress-induced release of mitochondrial double-stranded RNA activates RIG-I to drive osteogenic differentiation in aortic disease

**DOI:** 10.64898/2026.10.28.762541

**Authors:** Madeleine Graef, Felix Bender, Svenja Dudek, Paraskevi Vasileiadou, Sofía Soler, Thais Schlee-Guimaraes, Lisa Maria Steinheuer, Jasper Spitzer, Kevin Thurley, Radoslaw P. Nowak, Katharina Maser, Susanne Schmidt, Philip R Goody, Martin Schlee, Sebastian Zimmer, Georg Nickenig, Thomas Zillinger, Eva Bartok, Gunther Hartmann

**Affiliations:** Institute of Clinical Chemistry and Clinical Pharmacology, University Hospital, University of Bonn, Venusberg-Campus 1, 53127 Bonn, Germany; Institute for Experimental Haematology and Transfusion Medicine, University Hospital, University of Bonn, Venusberg-Campus 1, 53127 Bonn, Germany; Institute of Experimental Oncology, Biomathematics Division, University Hospital, University of Bonn, Venusberg-Campus 1, 53127 Bonn, Germany; Institute of Structural Biology, University Hospital, University of Bonn, Venusberg-Campus 1, 53127 Bonn, Germany; Heart Center Bonn, Molecular Cardiology, Department of Internal Medicine II, University Hospital Bonn, Venusberg-Campus 1, 53127 Bonn, Germany

**Keywords:** RIG-I, nucleic acid immunity, aortic valve stenosis, vascular calcification, mitochondrial RNA, JAK/STAT signaling, innate immunity

## Abstract

**Background:** Gain-of-function mutations in RIG-I and MDA5 (RIG-I-like receptors, RLR) cause Singleton-Merten syndrome, an interferonopathy with early aortic calcification. Here we set out to study a possible involvement of RIG-I-like receptor signaling in the formation of aortic valve stenosis in general. Methods and results: We investigated whether mechanical stress-induced RLR activation promotes osteogenic differentiation and calcification of primary human aortic cells, whether such RLR activation drives the development of aortic valve stenosis in vivo, and whether inhibition of downstream JAK/STAT activation ameliorates aortic valve stenosis. We found that activation of primary human aortic interstitial cells by exogenous RIG-I- and MDA5 agonists induced a type I interferon response and an osteogenic program including increased expression of BMP-2 and BSP2, and promoted calcification. These effects were RIG-I/MDA5 or MAVS-dependent, as assessed by CRISPR/Cas9 knockouts. The stress-granule protein G3BP1 and the Wnt receptor FZD5 were induced downstream of RLR activation and were required for osteogenic gene expression and calcification. To assess a potential endogenous source of RIG-I activation, mechanical stretch mimicking valvular tissue stress was applied in cell cuture, which induced a significant RIG-I-dependent interferon response. Sequencing of highly purified preparations of double-stranded RNA from stretched cells demonstrated a predominant mitochondrial origin of double-stranded RNA, and depleting mitochondrial RNA abolished the stretch-dependent response. The JAK/STAT inhibitor ruxolitinib suppressed the RIG-I-induced response including the expression of osteogenic factors *in vitro*. In a murine wire-injury model of aortic stenosis, MAVS deficiency attenuated valvular morphological changes and improved hemodynamic parameters. Ruxolitinib attenuated hemodynamic progression, prevented aortic stenosis development, and improved survival. Conclusions: Altogether, our results demonstrate that the mechanical-stress-induced release of mitochondrial double-stranded RNA activates RIG-I signaling in aortic interstitial cells, thereby driving osteogenic differentiation and calcification via a MAVS- and G3BP1-dependent pathway. JAK/STAT inhibition with ruxolitinib mitigates this process in a mouse model of aortic valve stenosis, identifying a candidate therapeutic strategy.

**Translational Perspective:** Aortic valve stenosis is a common and progressive disease, with no approved drug treatment. The only effective intervention is surgical or transcatheter valve replacement. This study shows that mechanical stress on human aortic interstitial cells leads to the leakage of mitochondrial double-stranded RNA, mistakenly recognized by the antiviral sensor RIG-I as foreign viral RNA, triggering a type I interferon response and osteogenic differentiation. Ruxolitinib, an oral Janus kinase inhibitor already approved for other conditions, reduced this response in cell culture and the development of disease in a mouse model of aortic valve stenosis. The results highlight the therapeutic potential of a ruxolitinib-based drug-repurposing strategy for aortic valve stenosis, a clinical strategy to be confirmed in clinical trials.

## Introduction

The innate immune system distinguishes pathogen-derived nucleic acids from host nucleic acids through a limited set of germline-encoded innate immune sensing receptors, among which the RIG-I-like receptors (RLRs) RIG-I (encoded by *DDX58*) and MDA5 (encoded by *IFIH1*) are the principal cytosolic immune sensors of double-stranded RNA (dsRNA) ^1–3^. RIG-I recognizes short, blunt-ended dsRNA bearing an uncapped 5′-triphosphate or 5′-diphosphate group, a structural signature generated as unavoidable by-product of viral and bacterial polymerase activity, whereas MDA5 preferentially detects longer dsRNA structures ^4–7^. Both RIG-I and MDA5 are broadly expressed in all somatic cells and immune cells, and converge on the common adaptor protein MAVS (mitochondrial antiviral-signaling protein) on the outer mitochondrial membrane, triggering IRF3 and NF-κB activation and a type I/III interferon (IFN) response ^2^.

Host RNA is normally shielded from RLR recognition. Self mRNA formed in the nucleus of eukaryotic cells undergo 5′-capping and 2′-O-methylation at the N1 position (cap1) before nuclear export. These modifications exclude nuclear mRNA from RIG-I binding. Mitochondrial RNA, however, is transcribed by a distinct polymerase (POLRMT) and does not undergo equivalent modifications and quality control. Through bidirectional transcription of the circular mt-DNA, mitochondria are a major source for endogenous dsRNA ^8,9^. When leaking into the cytoplasm, mitochondrial dsRNA can in principle be recognized by RIG-I as non-self ^10^. Notably, most mitochondrial dsRNA species carry a 5′-monophosphate rather than a 5′-triphosphate, which excludes RIG-I binding. The structural basis for this exclusion has recently been defined by our group ^11^. Failure of this molecular RIG-I escape mechanism, or increased availability of RLR ligands through insufficient clearance of self-nucleic acids or mitochondrial stress, have been shown to break self-tolerance to mt-RNA and lead to RLR-driven interferon responses ^9,12–14^.

Singleton-Merten syndrome (SMS) is a rare interferonopathy which is caused by monoallelic gain-of-function mutations in *DDX58* (RIG-I) or *IFIH1* (MDA5), which render the receptor constitutively active independent of exogenous ligand ^15–17^. Patients present with dental and skeletal anomalies together with early and often severe calcification of the aorta and aortic valve ^18,19^. SMS therefore demonstrates that constitutive RLR activation is sufficient to drive aortic and valvular calcification, providing genetic proof of concept for a possible involvement of RLR signaling in this pathology in general.

At the same time, the observation of aortic calcification in SMS raises an apparent paradox. Systemic type I induction and RLR activation through genuine viral infection or therapeutic administration of RLR agonists have not been associated with aortic valve calcification. Furthermore, decades of clinical experience with high-dose, long-term recombinant IFN-α therapy for chronic hepatitis B and IFN-β therapy for multiple sclerosis have not reported aortic or valvular calcification as adverse side effects, despite well-documented cardiovascular and hematological toxicity of these regimens ^20–22^. This dissociation suggests that systemic RLR/IFN pathway activation alone is not sufficient to calcify the aortic wall, and that aortic wall cells themselves may either be poorly responsive to circulating IFN, or that local, cell-intrinsic RLR activation, rather than a systemic type I interferon response is the relevant driver in SMS. So far, it has not been established whether resident aortic cells are capable of RLR activation, and what the consequences of such activation would be.

Aortic disease, including aortic valve stenosis (AVS), arises through active, cell-mediated osteogenic transdifferentiation of resident mesenchymal cells rather than passive mineral deposition^23,24^. Downregulation of contractile markers and upregulation of osteogenic transcription factors and matrix proteins (Runx2, osteopontin, bone sialoprotein, BMP-2) drive matrix-vesicle-mediated nucleation of hydroxyapatite within the vessel and valve wall ^25,26^. Toll-like receptors and other nucleic-acid-sensing PRRs have previously been implicated in this process in the aortic valve and vessel wall ^27–30^, but a specific role for the RLR/MAVS axis in aortic wall or valve cells outside the SMS context, and the identity of any physiological self-RNA ligand that might activate the RLR/MAVS axis, remain unresolved. The pathogenesis of AVS itself is still incompletely understood. Since the aortic valve is subject to substantial and cyclic mechanical loading during each cardiac cycle, and pathological deviations in wall stress and cyclic strain are established contributors to vascular remodeling and inflammation ^31–34^, we considered mechanical stress a plausible candidate trigger for local, ligand-dependent RLR activation in the aortic wall independent of any germline mutation like in SMS.

In this context, mechanical stress acting on cells of the aortic wall and valve may trigger spontaneous, self-RNA-driven activation of RIG-I, most plausibly through stress-associated release of mitochondrial dsRNA into the cytosol, thereby contributing to osteogenic transdifferentiation, calcification, and ultimately to the development of aortic disease. Downstream of RIG-I/MAVS, we considered the non-canonical Wnt pathway component G3BP1 (Ras-GTPase-activating protein SH3-domain-binding protein 1) as a candidate mechanistic link to osteogenic differentiation. G3BP1 is a core stress-granule nucleating protein with an established role in the translation of interferon-stimulated transcripts ^35^ and in cGAS-dependent DNA sensing ^36^, and has been directly implicated in RIG-I/MAVS-dependent, pro-sclerotic non-canonical Wnt signaling in SMS-mutant vascular smooth muscle cells ^37,38^. Finally, because RLR signaling converges on JAK/STAT-dependent interferon-stimulated gene transcription, and because the JAK1/2 inhibitor ruxolitinib has shown efficacy in interferonopathies mechanistically related to SMS, including STING-associated vasculopathy of infancy and SMS itself ^39,40^, pharmacological inhibition of the JAK/STAT pathway may limit RIG-I-driven osteogenic differentiation and the development of aortic valve stenosis in vivo.

Our results identify RIG-I and MAVS as functional drivers of osteogenic transdifferentiation and calcification in human aortic interstitial cells, implicate mechanical-stress-induced release of mitochondrial dsRNA as an endogenous trigger for this pathway, confirm a contribution of this axis to aortic valve stenosis in a wire-injury mouse model, and demonstrate that JAK/STAT inhibition with ruxolitinib limits stenosis progression. Together our results suggest a mechanistic link between the SMS paradigm of constitutive RLR activation and non-genetic, mechanically driven aortic disease.

## Methods

### Mice

Male C57/BL6-J wild type mice at the age of 10-12 weeks were obtained from Janvier Labs. Mavs^-/-^ mice were provided from the cooperating research group of Hiroki Kato at the Institute of Cardiovascular Immunology at the University Hospital Bonn. Rig-I^-/-^ mice were generated from a breeding program in our laboratory. Both knockouts were generated on a C57/BL6-J background. All mice were held in cages in groups of 4–5 animals under a 12 h light/dark cycle and at a constant temperature of 22 °C with ad libitum access to chow with or without ruxolitinib (2 g/kg) and drinking water. Control and treatment groups were kept separated at different cages. Only male mice were used because female mice have significantly lower trans-aortic peak blood flow velocity at baseline and display different hemodynamic changes following wire injury. All animal experiments were conducted in accordance with the Directive 2010/63/EU and the German Animal Welfare Act and approved by the local institutional animal care and use authorities (Landesamt für Verbraucherschutz und Ernährung Nordrhein-Westfalen, LAVE, approval number 81-02.04.2018.A250).

### Mouse model of aortic valve stenosis

Aortic valve stenosis was induced using a standardized wire injury-technique ^44,45^. The entire procedure, along with the relevant echocardiographic and MRT measurements in the mice, was carried out by trained technicians. Under echocardiographic guidance, a slightly bent angioplasty wire (ASAHI INTEC 0.36 mm diameter) was inserted through the right carotid artery and then advanced into the left ventricle. The wire was pushed into the apex and pulled back just below the aortic level ten times with subsequent 200 rotations at the level of the aortic valve (aortic valve injury group: AVS). The control groups had a sham operation in which the wire was inserted into the vessel without reaching and damaging the valve (control group without aortic valve injury: Sham). The surgical procedures were performed alternately on mice from the intervention and control groups. The surgeon was blinded to both the genotype and the treatment of the animals. Mice that did not survive the procedure were excluded from further analysis. No additional inclusion or exclusion criteria were defined. Surgery for the induction of aortic valve stenosis (AVS) was performed under intraperitoneal anesthesia with ketamine (100 mg/kg) and xylazine (16 mg/kg), dissolved in saline and administered at a maximum injection volume of 10 mL/kg. If required, anesthesia was prolonged by intraperitoneal administration of an additional 10–30% of the initial dose. Postoperative analgesia was provided with buprenorphine (0.1 mg/kg, subcutaneously), starting immediately after surgery and administered every 6 h during the light phase. During the dark phase, buprenorphine was provided via the drinking water (1 mL of buprenorphine solution, 0.3 mg/mL, diluted in 32 mL of drinking water). Analgesic treatment was maintained for at least 48 h postoperatively and extended when clinically indicated.

Echocardiography was performed in all mice two days before the wire injury for assessing the baseline levels of relevant parameters of aortic valve stenosis as previously described ^41^. After wire injury the mice had echocardiography every two weeks for a time frame of 6 weeks to monitor the development of aortic valve stenosis. The echocardiographic measurements included aortic valve peak blood flow velocity, ejection fraction (EF), valve orifice area, left ventricular (LV) function and volume, heart rate and wall thickness of the posterior wall. Echocardiography was performed under isoflurane anesthesia (induction: 3–5%; maintenance: 1–2%, by inhalation). Body temperature was maintained within the physiological range using a heating mat and monitored continuously with a rectal temperature probe. The measurements were performed with a Fijifilm Visualsonics Vevo 2100 and the analyses were produced using the Fijifilm Visualsonics VevoLab software. One day after the last echocardiographic measurement (42 days post wire injury), mice were sacrificed and organs were collected for further examination. The whole heart was snap-frozen in Tissue-Tek O.C.T medium for histological analysis of the aortic valve. From each heart approximately 10 slides, with 6–8 cryosections of aortic valve tissue, each starting from the aortic root, were generated using a Leica microtome. Euthanasia was performed by exsanguination under general anesthesia induced with ketamine (100 mg/kg) and xylazine (16 mg/kg), administered intraperitoneally as described above. Following midline laparotomy, the caudal vena cava was exposed and punctured, and approximately 1 mL of venous blood was collected, resulting in euthanasia by exsanguination. Subsequently, the thoracic cavity was opened, and the circulatory system was perfused with cold saline via the left ventricle.

### Histological analysis of mouse aortic valve tissue

#### Immunofluorescence staining

Slides with sections of mouse aortic valve tissue were taken out from -80°C and left air-dry for approximately 15–20 minutes. Firstly, a fixation was performed by immersion of the slides into ice-cold acetone for 20 min, followed by three washes with 1x PBS, each for 5 min. Sections were incubated with blocking buffer (1x PBS with 2% BSA) for 1 hour at RT. After three washes with 1x PBS, 5 min each, the sections were incubated overnight at 4 °C with the primary antibody CD45 (1:100) or CD68 (1:100) diluted in immunofluorescence blocking buffer (1% BSA and 2% normal goat serum in PBS). Following this, the sections were washed three times with 1x PBS, each for 5 min, and then incubated with the secondary antibody goat anti-rat Alexa Fluor 555 (1:500) diluted in blocking buffer for 1 hour at RT in the dark. After three more washes with 1x PBS, each for 5 min, the sections were stained with DAPI (300nM in PBS) for 10 min at RT in the dark. Finally, the sections were washed three times with 1x PBS for 5 min each and mounted with ProLong Gold Antifade Mountant. Images were acquired with a Leica DMi8 epifluorescence microscope and a Leica DFC9000 GT camera.

#### Hematoxylin and eosin (H&E) staining

Aortic valve sections were H&E-stained by the *Histology Core Facility* of the Medical Faculty at the University of Bonn.

### Immunofluorescence staining of cultured cells

1.5×10^4^ HAoSMCs per well were seeded in a 96-well PhenoPlate (PerkinElmer) and stimulated for 72 h as described above. Cells were fixed with buffered 4% paraformaldehyde (PFA) solution for 20 min at 4°C, permeabilized with 0.1% Triton X-100 in PBS for 10 min, and blocked with immunofluorescence blocking buffer for 30 min at room temperature. Cells were incubated overnight at 4 °C with a primary antibody against G3BP1 diluted in blocking buffer. After washing, cells were incubated with Alexa Fluor 647-conjugated donkey anti-rabbit secondary antibody (1:500) for 1 h at room temperature. Nuclei were counterstained with DAPI (1:1000) for 20 min. Fluorescence images were acquired using a Leica DMi8 epifluorescence microscope equipped with a Leica DFC9000 GT camera and images were processed with Fiji (ImageJ).

### Flow cytometry of murine peripheral blood

Peripheral blood was collected from the submandibular vein 3 days after wire-injury in heparinized tubes and diluted with PBS. Following centrifugation (400 × g, 4 min), erythrocytes were lysed using ammonium chloride-based lysis buffer for 5 min. Lysis was stopped by addition of PBS, and cells were washed by centrifugation (400 × g, 4 min, 4°C). Cell suspensions were transferred to 96-well U-bottom plates and stained with Zombie Aqua viability dye (1:2000) for 30 min at room temperature in the dark. After washing, Fc receptors were blocked (Fc Block, 1:200, 5 min), followed by staining with fluorochrome-conjugated antibodies against CD45, CD3, CD11c and NK1.1 for 30 min on ice in the dark. Cells were washed, fixed using Low Fix Buffer and stored at 4°C until acquisition. Flow cytometry data were acquired on a 5-Laser Sony ID700™ Spectral Analyzer and evaluated using FlowJo 11 (BD Biosciences). Cells were sequentially gated on singlets, viable cells, and CD45⁺ leukocytes before analysis of immune cell subsets. T cells were identified as CD3⁺ cells, dendritic cells as CD11c⁺ cells and NK cells as NK1.1⁺CD3⁻ cells. Immune cell populations are presented as the percentage of CD45⁺ leukocytes.

### Aortic cell culture and stimulation

Human aortic valve tissue was obtained from patients undergoing aortic valve replacement at the Heart Center Bonn, University Hospital Bonn. The clinical study was approved by the Ethics Committee of the University of Bonn (approval number 078/17). All participants provided written informed consent. The study was conducted in accordance with the Declaration of Helsinki and Good Clinical Practice guidelines. Following excision, valve cusps were incubated in 600 U/ml collagenase II in VIC primary medium (DMEM supplemented with 10% FBS, 1% penicillin/streptomycin, and 44 mM NaHCO₃) for 10 min at 37 °C with gentle agitation to first remove the endothelial cell layer. The remaining valve tissue was then minced and further digested in the same collagenase II solution for 20 h at 37 °C with gentle agitation. The resulting cell suspension was filtered (100 µm) and centrifuged twice (1,000 rpm, 15 min), and VICs were seeded on collagen-coated (5 µg/cm²) T75 flasks. Cells were cultured in VIC medium (DMEM, 10% FBS, 1% penicillin/streptomycin) at 37 °C, 5% CO₂, 95% humidity, and experiments were performed at passage 5–6. Human aortic smooth muscle cells (HAoSMC) were purchased from PromoCell. Cells were cultured in Smooth Muscle Cell Growth Medium 2 (PromoCell) with 1x Antibiotic-Antimycotic (LifeTechnologies) until passage 11. Cells were cultivated at 37 °C in a humified atmosphere with 5 % CO_2_ and the medium was changed every 2–3 days. Cells were seeded when they reached 80–90 % confluence. 24 h after seeding, cells were transfected with RIG-I ligand IVT4 (5’-triphosphate hairpin RNA: 3pRNA, 400 ng/ml) ^42^, total RNA from EMCV-infected Vero E6 cells (800 ng/ml) or a negative control RNA (3p-ssRNA, 400 ng/ml) using Lipofectamine 2000 (Thermo Fishe Scientific), or treated with recombinant human IFNβ (Peprotech, 1000 U/ml) for 72 h if not indicated otherwise. Calcification was induced in pro-calcifying medium (PCM; Smooth Muscle Cell Growth Medium 2 containing 1x Antibiotc-Antimycotic, 50 µg/ml ascorbic acid and 3, 4 or 5 mM disodium hydrogen phosphate). To inhibit the JAK-STAT pathway in HAoSMCs, cells were preincubated with 6 µM Ruxolitinib for 45 min before stimulation. DMSO was used as control.

### Application of cyclic mechanical stretch

To apply mechanical stress to HAoSMCs, cells were plated in a type I collagen-coated BioFlex® 6-well culture plate (1.8 × 10^5^ cells/well). After 24 h cells were stimulated with or without additional cyclic stretch. Cyclic stretch was applied to the culture plates at 0.5 Hz and 15% elongation by using a Flexcell® Tension System (FX-6000T) inside a 5% CO_2_ incubator at 37 °C for 72 h. Static cultures served as control. To inhibit mitochondrial RNA transcription, HAoSMCs were preincubated with the mitochondrial RNA polymerase inhibitor IMT1 (10 µM) for 72 h prior to application of mechanical stretch. DMSO-treated cells served as vehicle controls.

### Generation of polyclonal knockout cells by CRISPR/Cas9

Polyclonal knockouts were generated via the Alt-R CRISPR-Cas9 system (IDT). crRNAs targeting G3BP1, FZD5, RIG-I, MDA5 and MAVS were designed using the Alt-R HDR design tool (IDT). A non-targeting crRNA was used as control. Delivery into cells was performed using the Neon Transfection System (Thermo Fisher Scientific) following the Alt-R CRISPR-Cas9 user guide. Briefly, crRNA and tracrRNA were annealed at 44 µM and 22 pmol of the crRNA:tracrRNA duplex were mixed with 18 pmol Alt-R Cas9 enzyme to form the RNP complex. 1.8 × 10^5^ HAoSMCs were mixed with the Cas9 RNP (1.8 µM crRNA:tracRNA, 1.5 µM Cas9) and 1.8 µM Alt-R Cas9 electroporation enhancer in Buffer R and were electroporated using the Neon Transfection 10 µL kit (1475 V, 2 pulses, 20 ms). After electroppration, cells were transferred into a 12-well plate containing 1 ml pre-warmed medium per well. Cells were incubated as described above and expanded for subsequent experiments. KO efficiency was determined by Western blotting and/or Sanger sequencing of PCR-amplified genomic loci. For Sanger sequencing, genomic loci were amplified with DreamTaq Polymerase and purified PCR products were sequenced by Microsynth (Goettingen, Germany). Frequency of insertions and deletions (INDELs) was quantified from ab1 files using TIDE^43^. Cells were considered as polyclonal KO when at least 70 % of the sequences showed INDELs.

### RNA isolation and qRT-PCR

RNA was isolated using the RNeasy mini kit (Qiagen) modified as described in ^46^. Reverse transcription was performed using random hexamers (Integrated DNA Technologies) and RevertAid reverse transcriptase (Thermo Fisher Scientific) according to manufacturer’s instructions. cDNA was subjected to real-time quantitative PCR (qPCR) using the EvaGreen QPCR-mix II (ROX) (Biobudget) on a 384-well QuantStudio 5 machine (Thermo Fisher Scientific). Ct values were normalized to *GAPDH* or HPRT, as indicated. Primers are listed in Table 1.

**Table 1:**
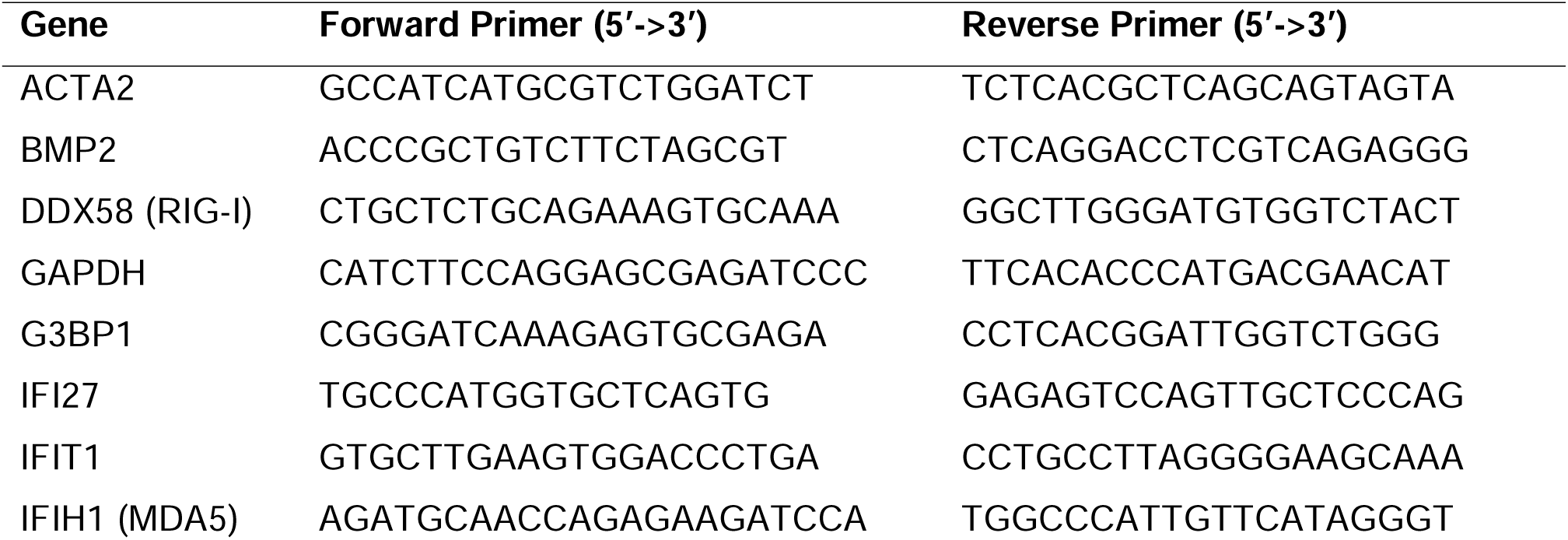
qPCR primers.

### 3’-mRNA sequencing

RNA sequencing libraries were prepared using the QuantSeq 3′ mRNA-Seq Library Prep Kit FWD (Lexogen) according to the manufacturer’s instructions and sequenced on an Illumina HiSeq 1500 platform at the Next Generation Sequencing Core Facility of the University Hospital Bonn. Reads were aligned to the human reference genome (hg19) using STAR (v2.6.1d), and transcript abundance was quantified with the Partek E/M algorithm. Differential gene expression analysis was performed using DESeq2 (v1.40.2) ^44^ and log₂ fold changes were shrunk using the *ashr* method ^45^. Gene Ontology (GO) pathway enrichment analysis was performed using clusterProfiler (v4.8.3) ^46^. Data processing and visualization were carried out using the tidyverse package collection (v2.0.0) ^47^.

### Selective sequencing of double-stranded RNA (SALDOR-Seq)

Cellular dsRNA was sequenced by SALDOR-Seq as described here ^48^. Briefly, total RNA from untreated HAoSMCs was isolated with TRIzol and digested with RNases A/T1 and DNase I to isolate dsRNA. Purified dsRNA was dephosphorylated, denatured, adapter-ligated and reverse-transcribed using an RT primer bearing a 5’ overhang with the Oxford Nanopore cDNA-PCR primer binding site. cDNA libraries were barcoded, amplified by single-primer PCR and prepared for sequencing using the Oxford Nanopore Technologies cDNA-PCR Barcoding Kit (V14). Sequencing was carried out on a MinION Mk1B device equipped with MinION or Flongle flow cells.

### Western blot analysis

Cell lysates were prepared in Laemmli buffer and proteins were size-separated via sodium dodecyl-sulfate (SDS) polyacrylamide gel-electrophoresis (PAGE) according to standard procedures. Subsequently, proteins were transferred onto a 0.45 µm nitrocellulose membrane via wet-transfer for 90 min at 100 V using a Mini Trans-Blot® Cell (Bio-Rad). Afterwards, the membrane was blocked in blocking buffer (5% BSA in TBS-T) for 1 h at RT. The membrane was washed with TBS-T and then incubated with the primary antibody diluted in blocking buffer overnight at 4 °C. Next, membrane was washed and incubated with fluorophore-conjugated secondary antibodies diluted 1:10,000 in blocking buffer at RT for 1.5 h. After washing, fluorescence was detected using an Odyssey Fc Imager (Licor).

### Alizarin red S staining

Cells were fixed for 20 min at 4 °C in buffered 4% PFA solution (Merck). After two washes with distilled water, cells were incubated with 2% alizarin red S solution, pH 4.2 (Sigma Aldrich) for 30–45 min. Excess stain was removed with distilled water. Alizarin red stain was quantified by extraction with hexadecyl pyridium chloride and the absorbance was measured at 550 nm.

### BLaER1 cell differentiation and stimulation

BLaER1 cells were kindly provided by Thomas Graf, Centre for Genomic Regulation, Barcelona, Spain. Cells were cultured in RPMI 1640 medium supplemented with 10 % FCS and 100 U/ml penicillin/streptomycin. For differentiation to monocyte-like cells, BLaER1 cells were seeded at 7×10^4^ per 96-well and kept for five days in media containing 100 nM 17-β -estradiol (Sigma-Aldrich), 10 ng/ml IL-3 (Thermo Fisher Scientific) and 10 ng/ml M-CSF (Thermo Fisher Scientific). Cells were stimulated with 0.8 µg/ml RNA or DNA using 2 µl/ml Lipofectamine 2000 (Thermo Fisher Scientific). After 24 h, IFNα concentration in supernatants was measured by ELISA.

### Measurement of secreted cytokines

IP-10 (CXCL10) levels in cell culture supernatants were determined using the BD OptEIA™ Human IP-10 ELISA Set (BD Biosciences). IFNα was measured using a human IFNα ELISA kit (Thermo Fisher Scientific, BMS216MST). Type I IFN bioactivity was measured using HEK-Blue IFN-α/β reporter cells. For that, HEK-Blue cells were seeded at 5×10^4^ per 96-well and incubated with medium containing 10% supernatant from the stimulated cells. After 24 h, medium was incubated with pNPP substrate for 30 min at 37 °C. Reaction was stopped with 30 mM EDTA and absorbance at 405 nm was measured with a Cytation 3 Cell Imaging Multi-Mode Reader. Type I IFN activity in the supernatant was calculated from a standard curve with recombinant IFN-β and is expressed as U/ml.

## Statistical analysis

All results are shown as mean ± SEM. Statistical significance was determined with GraphPad Prism 10 using, paired t-test, one-way ANOVA with Dunnett’s or Tukey’s multiple comparisons test or two-way ANOVA with Dunnett’s, Tukey’s, or Sidak’s multiple comparisons test and a confidence interval of 95%. Survival analysis was performed using a two-sided log-rank test. Significances are indicated as followed: * (p < 0.05), **(p < 0.01), *** (p < 0.001), **** (p ≤ 0.0001), ns: not significant.

## Results

### RIG-I-like receptor activation induces a type I interferon response and osteogenic transcriptional program in human aortic cells

To determine whether primary human aortic cells (valve interstitial cells, VICs; primary human smooth muscle cells, HAoSMCs) are capable to sense and respond to RIG-I-like receptor (RLR) ligands, we first assessed cytokine secretion, interferon-stimulated gene (ISG) induction, and the downstream transcriptional consequences of RLR activation in these cells. Stimulation of primary human aortic cells with the RIG-I agonist 3pRNA significantly increased secretion of the ISGs compared with medium control (Fig. 1A and suppl. Fig. 1). Induction of ISGs (*IFIT1*, *IFI27*, *RIG-I*, *MDA5)* following stimulation with IFNβ or 3pRNA was time-dependent (Fig. 1B). Transcriptomic analysis (3′mRNA sequencing) of 3pRNA-versus control RNA-stimulated cells confirmed that RIG-I activation induces a broad interferon-associated transcriptional signature, including upregulation of the ISGs *IFITM1* and *ISG15*, but also of the osteogenic transcription factors *SOX8* and *CITED1*, alongside downregulation of the calcification inhibitor *MGP* (Fig. 1C). Gene Ontology enrichment analysis of this dataset identified significant enrichment for terms directly relevant to aortic calcification, including negative regulation of bone remodeling, osteoclast differentiation, positive regulation of bone mineralization, and calcium ion homeostasis (Fig. 1D). Together, these results show that RLR activation in primary human aortic interstitial cells triggers a type I interferon response while simultaneously engaging a transcriptional program associated with osteogenic differentiation, identifying RLR signaling as a candidate molecular link between innate immune activation and vascular calcification. The osteogenic markers *BMP-2* and *BSP2* were significantly induced by the RIG-I ligand IVT4 (3pRNA) and by RNA from EMCV-infected cells known to activate MDA5, but not by IFNβ alone, with the strongest induction at 48–72 h (suppl. Fig. 2A). This suggests a cell-intrinsic mechanism directly mediated through RLR signaling rather than through secreted type I IFNs. To further unravel the involvement of the components of the RLR signaling pathway, we used CRISPR/Cas9 to generate polyclonal HAoSMCs deficient in RIG-I, MDA5 or MAVS via transient Cas9 RNP delivery. Functional disruption was confirmed by reduction of IP-10 secretion in cells transfected gRNAs targeting RIG-I, MDA5 or MAVS compared to a non-targeting gRNA (ctrl. gRNA) following RIG-I or MDA5 stimulation, respectively (Fig. suppl. Fig. 2B). Under EMCV-RNA stimulation, *BMP-2* mRNA induction was reduced in MDA5- and in MAVS-deficient cells relative to gRNA control, whereas IFNβ- and control-RNA-stimulated conditions showed no genotype-dependent differences (suppl. Fig. 2C). We next hypothesized that osteogenic differentiation following RLR activation may lead to calcification in our primary cell model. We therefore performed functional calcification assays (Alizarin Red staining after 7 days in pro-calcifying medium) (Fig. 2A), which revealed that calcification increased with NaH₂PO₄ dose and was most pronounced under 3pRNA and EMCV-RNA stimulation at 5 mM NaH₂PO₄ (Fig. 2B). In MAVS-deficient cells, Alizarin Red staining was visibly reduced relative to control gRNA under 3pRNA and EMCV-RNA stimulation at 4–5 mM NaH₂PO₄ (Fig. 2C). Together, RIG-I- and MDA5-mediated signaling through MAVS drives osteogenic gene expression with statistically robust, pathway-dependent effects at the cytokine and mRNA level. The corresponding reduction in calcification in MAVS-deficient cells is consistent with this model.

**Figure 1:**
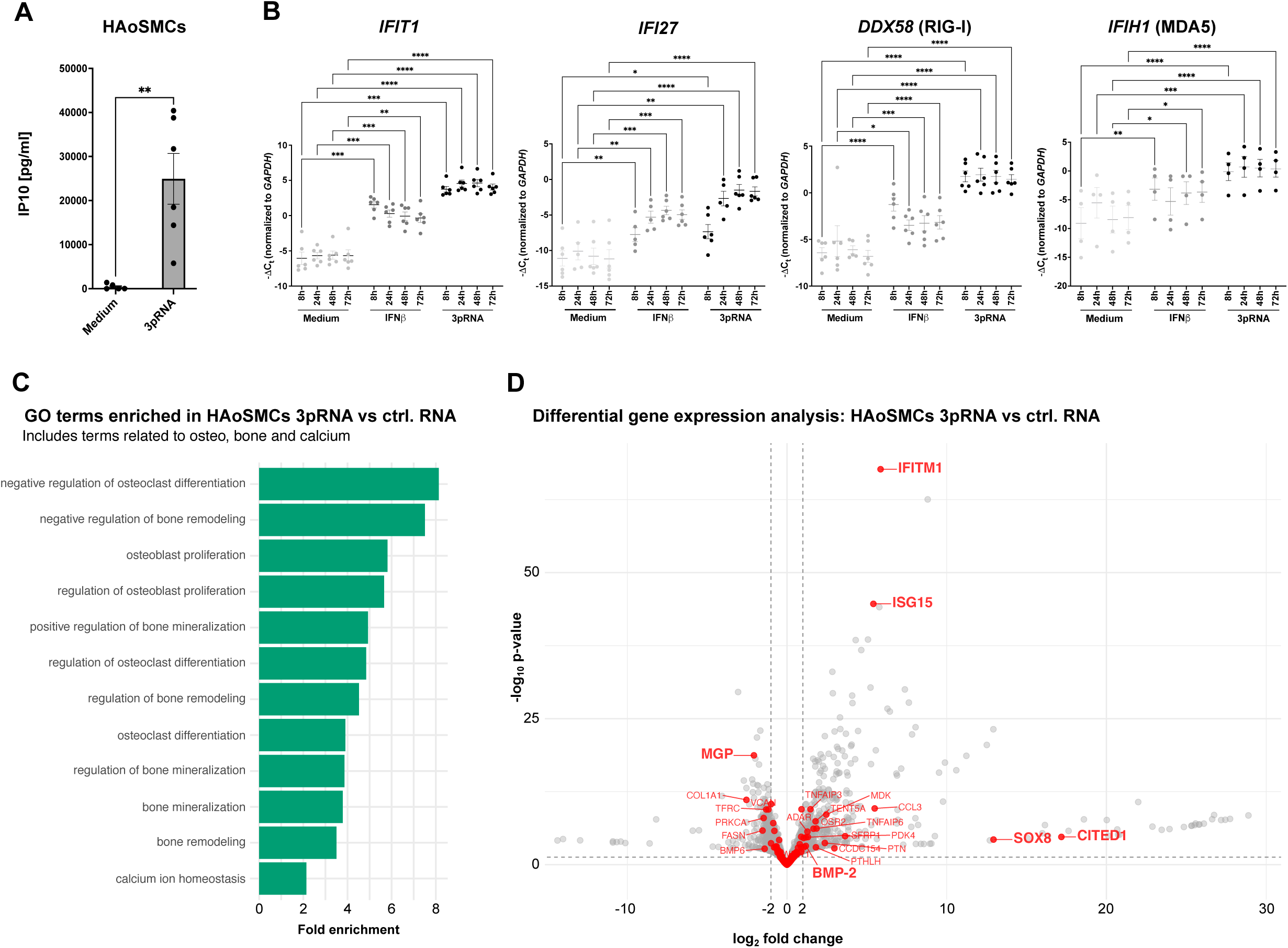
RIG-I activation of human aortic smooth muscle cells (HAoSMCs) induces the expression of interferon-stimulated genes including RIG-I and MDA5, and of gene sets associated with osteogenic differentiation, bone remodeling and regulation of mineralization. HAoSMCs were stimulated with 3pRNA (IVT4, RIG-I agonist) or IFNβ. (A) After 24 h, IP-10 was measured in the supernatants by ELISA (n=6 donors). (B) Expression of the interferon-stimulated genes *IFIT1*, *IFI27*, *DDX58* (RIG-I) and *IFIH1* (MDA5) (both ISGs) was analysed by qRT-PCR (n=6 donors). (C, D) Transcriptomic analysis and gene enrichment analysis were performed after 3’mRNA sequencing (n=3 independent experiments). Data are shown as mean ± SEM. * p<0.05, ** p<0.01, *** p<0.001, **** p<0.0001, ns = not significant; statistical significance was calculated by one-way ANOVA with Dunnett’s multiple comparison test (A) or two-way ANOVA with Dunnett’s multiple comparison test (B, C). Significance only shown for the comparisons between the different gRNA conditions.

**Figure 2.**
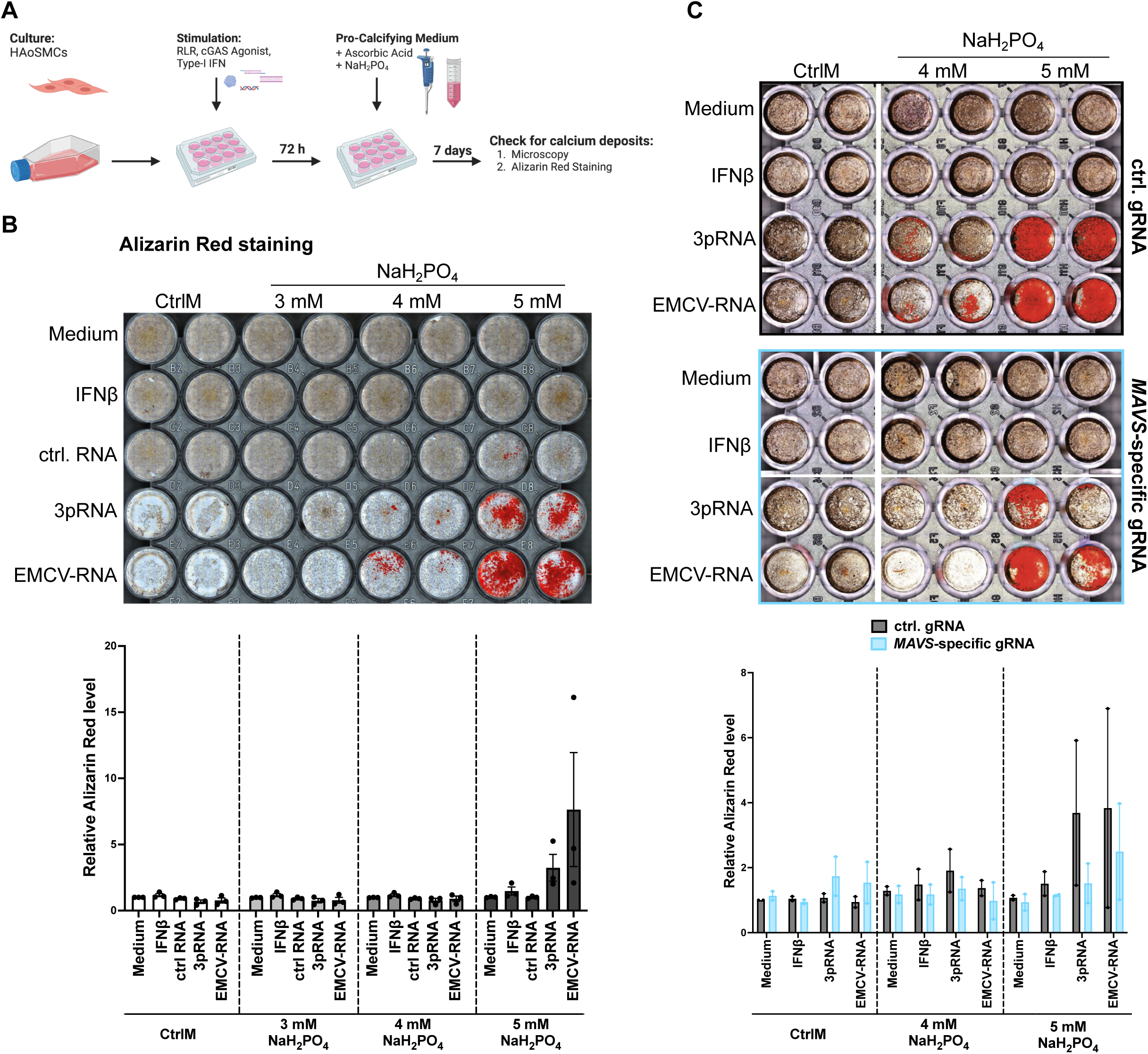
Activation of RIG-I-like receptors induces MAVS dependent calcification in human aortic smooth muscle cells. HAoSMC were incubated in the presence of RIG-I agonist (3pRNA), MDA5 agonist (EMCV-RNA), control RNA (ctrl RNA), IFNβ or medium. After 72 hours, pro-calcifying medium containing 50 µg ascorbic acid and 3, 4 or 5 mM NaH_2_PO_4_, or control medium were added. After 7 days, calcium deposition was quantified by Alizarin Red staining. Alizarin Red was dissolved in hexadecylpyridium chloride (CPC) and the absorbance was measured at 550 nm. A graphical overview is provided in (A). (B) The upper image shows a representative Alzarin Red staining of a cell culture plate at day 7 is shown. The graph below shows the means±SEM from primary HAoSMC from three different donors. (C) RLR-induced calcification at day 7 in MAVS-deficient primary HAoSMC (MAVS-targeting gRNA, lower panel) compared to MAVS-competent control HAoSMC (ctrl. gRNA, upper panel). The bottom graph shows the means±SEM of quantitative calcification from primary HAoSMC from two different donors.

### RIG-I-like receptor-driven osteogenic differentiation and calcification require G3BP1

We next investigated the RLR-associated stress-granule protein G3BP1 and the Wnt receptor FZD5 as candidate downstream effectors linking RLR activation to osteogenic differentiation and calcification. *G3BP1* and *FZD5* mRNA were both significantly induced by the RIG-I ligand 3pRNA and the MDA5 ligand EMCV-RNA, but not by IFNβ alone, reaching significance from 48 h onward (Fig. 3A). Immunofluorescence confirmed increased G3BP1 protein staining following 3pRNA stimulation relative to medium or IFNβ controls (Fig. 3B). 3pRNA-induced *G3BP1* mRNA induction was significantly reduced in RIG-I-deficient cells relative to control gRNA-transfected cells (Fig. 3C), indicating that G3BP1 induction downstream of RLR activation is RIG-I-dependent. In the calcification assay, both G3BP1-knockout and FZD5-knockout cells showed reduced Alizarin Red staining compared to wild-type control cells (ctrl. gRNA) stimulated with 3pRNA and EMCV-RNA (suppl. Fig. 3A). Consistent with a requirement for G3BP1, 3pRNA-induced *BMP-2* mRNA induction was significantly reduced specifically in G3BP1-but not in FZD5-deficient cells relative to gRNA control, while cytokine secretion was not affected (suppl. Fig. 3B). Together, these results position G3BP1 as a RIG-I-dependent downstream node that is required for RLR-driven osteogenic gene expression and calcification.

**Figure 3:**
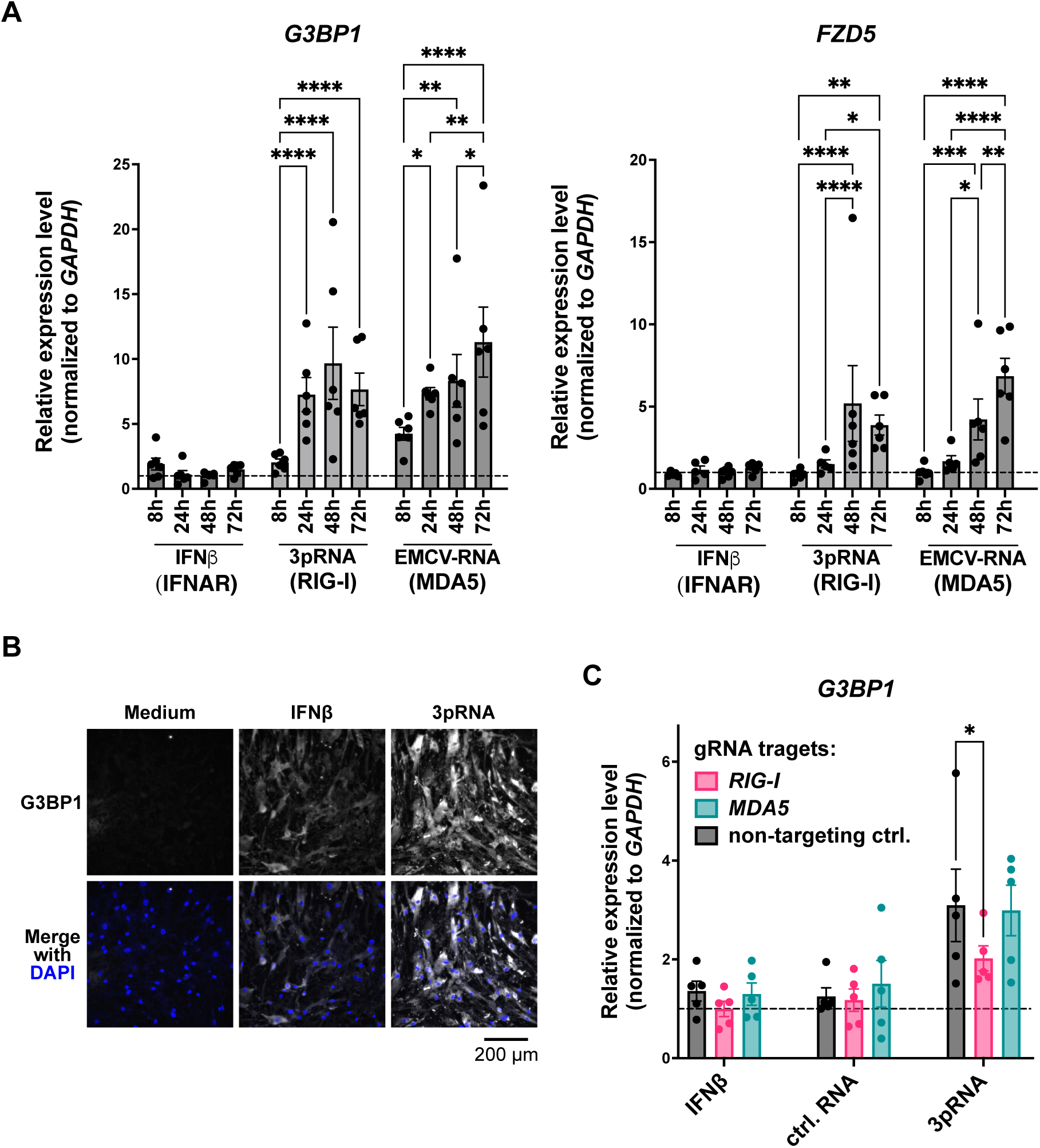
G3BP1 and FZD5 are upregulated by RIG-I-like receptor activation. Primary HAoSMCs were stimulated with IFNβ, 3pRNA (RIG-I) or EMCV-RNA (MDA5). (A) The mRNA expression of *G3BP1* (stress granule, left graph) and *FZD5* (Wnt receptor, right graph) were analyzed (n=6). (B) G3BP1 protein expression in response to 3pRNA (RIG-I) compared to IFNβ and medium control was analyzed by immunofluorescence staining (72 h, scale bar indicates 200 µm). (C) Induction of G3BP1 mRNA by the RIG-I agonist 3pRNA (versus IFNβ and control RNA) was quantified in RIG-I-deficient, MDA5-deficient and control HAoSMCs (72 h, n=5). Data are shown as mean ± SEM. * p<0.05, ** p<0.01, *** p<0.001, **** p<0.0001, ns = not significant; statistical significance was calculated by two-way ANOVA with Tukey’s multiple comparison test (A) or two-way ANOVA with Dunnett’s multiple comparison test (C).

### Mechanical stress is sufficient to activate type I interferon signaling in human aortic cells via RIG-I and MAVS

To test whether the RLR pathway contributes to disease progression *in vivo*, we used the wire-injury model of aortic valve stenosis in wild-type (WT), *Rig-i*⁻^/^⁻ and *Mavs*⁻^/^⁻ mice, with echocardiographic, histological, and immunofluorescence assessment over a 42-day time course (suppl. Fig. 4A). By echocardiography, peak transvalvular flow velocity (hemodynamic parameter positively correlated with the extent of aortic valve stenosis) was significantly lower in MAVS-deficient mice than in WT at 28 and 42 days post-injury (dpi) (suppl. Fig. 4B); RIG-I-deficient mice showed a decreasing trend at all timepoints. Aortic valve thickness at 42 dpi was reduced (trend) for *Rig-i*⁻^/^⁻ mice and *Mavs*⁻^/^⁻ mice (suppl. Fig. 4C). Infiltration of CD45-(leukocytes) and CD68-(myeloid immune cells) positive cells likewise was lower (trend) for *Rig-i*⁻^/^⁻ mice and *Mavs*⁻^/^⁻ mice (suppl. Fig. 4D). MAVS deficiency showed a significantly reduced hemodynamic disease progression after wire injury, consistent with a functional contribution of constitutive RLR signaling to aortic disease *in vivo*.

Because aortic cells are constantly exposed to mechanical stress, and even more so in disease, we tested whether cyclic mechanical stretch is sufficient to activate RLR-dependent type I IFN signaling, independent of exogenous ligand. Cyclic stretch (FlexCell Tension System, 0.5 Hz, 15% elongation) significantly increased IFNβ secretion at 72 h, with no significant change at earlier timepoints (Fig. 4A). At the mRNA level, *IFIT1* reached significance at 72 h, and *RIG-I* mRNA (RIG-I is an ISG) was significantly increased at 48 h and 72 h (Fig. 4B). Combining stretch with the RIG-I ligand 3pRNA did not produce a further significant increase in IFNβ protein secretion over either medium or 3pRNA alone without stretch (Fig. 4C). For *IFIT1* and *RIG-I* mRNA, stretch produced a significant increase in medium-treated cells but not in 3pRNA-treated cells, consistent with a ceiling effect from the already-maximal 3pRNA stimulus (Fig. 4D). Stretch-induced *IFIT1* upregulation was significantly blunted in RIG-I- and MAVS-deficient cells relative to wild-type cells (ctrl gRNA) (Fig. 4E). Next, total RNA was extracted from stretched or unstretched human aortic interstitial cells and transfected into the human monocyte-like BLaER1 cell line which is an established system for studying RLR signalling. Total RNA from stretched cells triggered significantly higher IFNα secretion in a RIG-I- and MAVS dependent manner (Fig. 4F, n = 4 donors, 3pRNA and HT-DNA used as positive controls). Together, these data demonstrate that mechanical stress, like present in the aorta, can induce the expression of endogenous RIG-I agonists triggering a type I IFN response.

**Figure 4:**
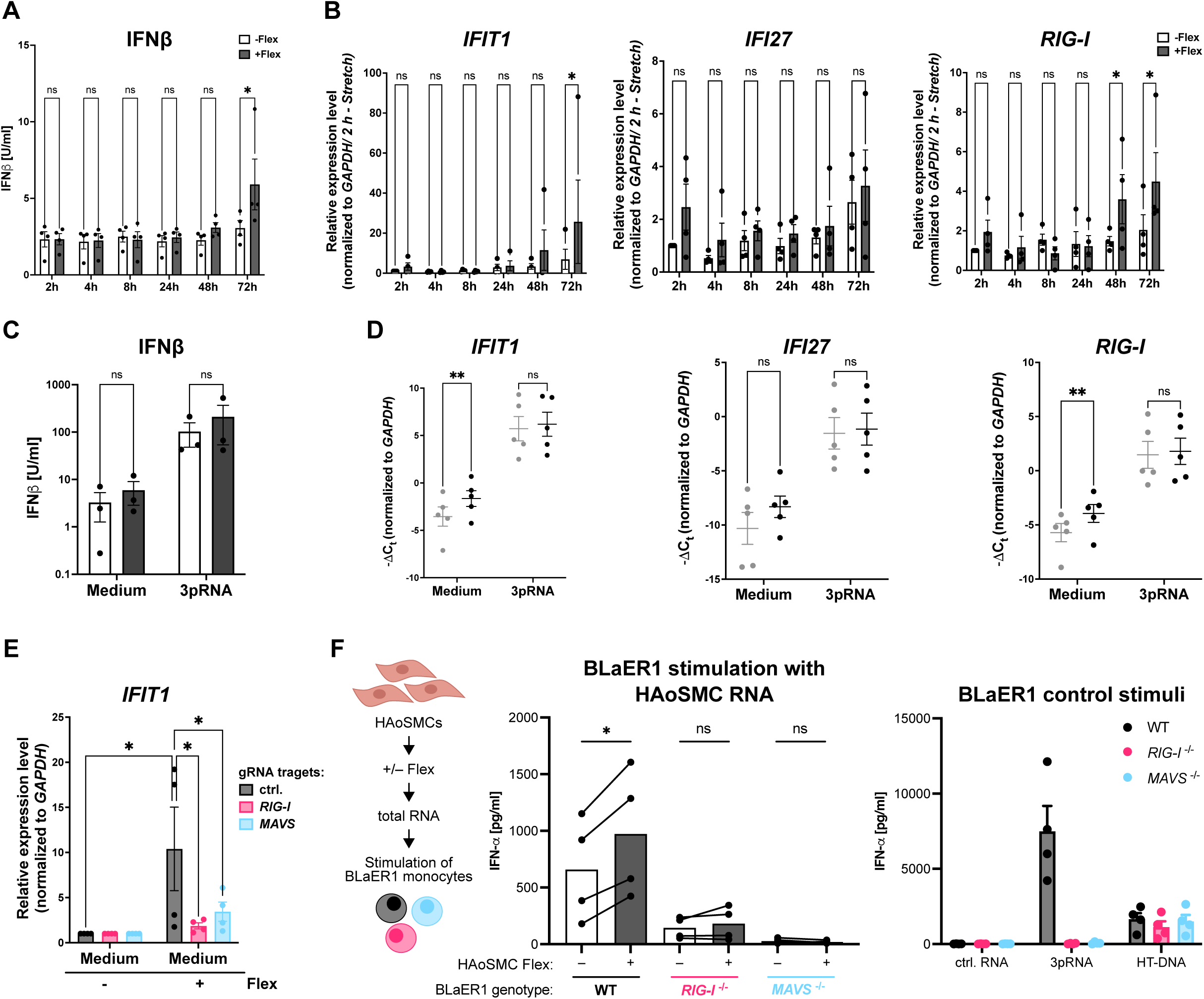
Stimulation of type I IFN and of IFN-inducible genes (ISG) by mechanical stress is RIG-I- and MAVS-dependent. HAoSMCs were exposed to cyclic mechanical stretch (+Flex) using a FlexCell Tension System (0.5 Hz, 15 % elongation, 72 h). (A) Kinetics of mechanical stretch-induced IFNβ secretion (n=3 donors). (B) Kinetics of mechanical stretch-induced expression of the interferon-stimulated genes *IFIT1*, *IFI27* and *RIG-I* (n=4 donors). (C, D) HAoSMCs were exposed to cyclic mechanical stretch in the presence or absence of the RIG-I agonist 3pRNA. After 72 hours, IFNβ (n=3) (C) and *IFIT1*, *IFI27* and *RIG-I* (D) were analyzed (n=5 experiments with 5 different donors). (E) RIG-I- and MAVS-deficient HAoSMCs showed reduced stretch-induced expression of the interferon-inducible gene *IFIT1* compared to control HAoSMCs (n=4 donors). (F) Total RNA isolated from stretched or non-stretched HAoSMCs (n=4 donors) was transfected into BLaER1 cells of indicated genotype. Stimulation with control stimuli is shown on the right. Data are shown as mean ± SEM. (* p<0.05, ** p<0.01, *** p<0.001, **** p<0.0001, ns = not significant; two-way ANOVA with Sidak’s multiple comparison test (A–E); two-tailed paired t-test (F).

To identify the endogenous ligand responsible for stretch-induced RIG-I activation, we characterized the origin of endogenous double-stranded RNA (dsRNA) species in human aortic interstitial cells using targeted dsRNA-sequencing via SALDOR-Seq ^48^. In line with previous studies on cellular dsRNA from other cell-types ^9,12–14^ the vast majority of endogenous dsRNA in aortic cells mapped to the mitochondrial chromosome (suppl. Fig. 5A). Due to the striking dominance of mitochondrial dsRNA (mt-dsRNA), we used the inhibitor of mitochondrial transcription 1 (IMT1) to deplete mtRNA and study a possible role of mt-dsRNA in the mechanical stress-induced RIG-I activation. As expected, IMT1-treatment significantly reduced levels of the constitutive single-stranded mitochondrial mRNA transcripts *COX1* and *ND1* with no difference with and without mechanical stress (±Flex) (suppl. Fig. 5B). We then checked whether IMT1 treatment affected the induction of ISGs through mechanical stress. *IFIT1*, *IFI27 and ISG15* mRNAs were increased through mechanical stress, an effect which was ablated in IMT1-treated human aortic cells (suppl. Fig. 5C, n = 5 donors). Together, these results suggest that mt-dsRNA is the predominant endogenous dsRNA species detectable, and its pharmacological depletion is associated with loss of the stretch-induced ISG response, supporting a model in which mechanical stress drives RIG-I activation through the release of mt-dsRNA.

### The JAK/STAT inhibitor ruxolitinib suppresses RIG-I-dependent osteogenic differentiation in vitro and ameliorates wire-injury-induced aortic valve stenosis *in vivo*

Finally, we tested whether pharmacological JAK/STAT inhibition with ruxolitinib, acting downstream of RLR signaling, is capable of reducing RLR-driven type I interferon- and osteogenic responses *in vitro* and disease progression *in vivo*. Ruxolitinib pretreatment of aortic interstitial cells significantly reduced IP-10 secretion induced by 3pRNA and by EMCV-RNA relative to DMSO control, with no significant effect in the control-RNA or medium conditions (Fig. 5A). In the presence of ruxolitinib, stimulation with IFNβ, 3pRNA, and EMCV-RNA showed reduced levels of RIG-I and MAVS protein (Fig. 5B). At the mRNA level, ruxolitinib significantly reduced *IFIT1* induction upon IFNβ, 3pRNA, and EMCV-RNA stimulation relative to DMSO (Fig. 5C), and significantly reduced *BMP-2* induction upon stimulation with 3pRNA and EMCV-RNA stimulation (Fig. 5D). Ruxolitinib also reduced calcification in the Alizarin Red assay (results not shown).

**Fig. 5:**
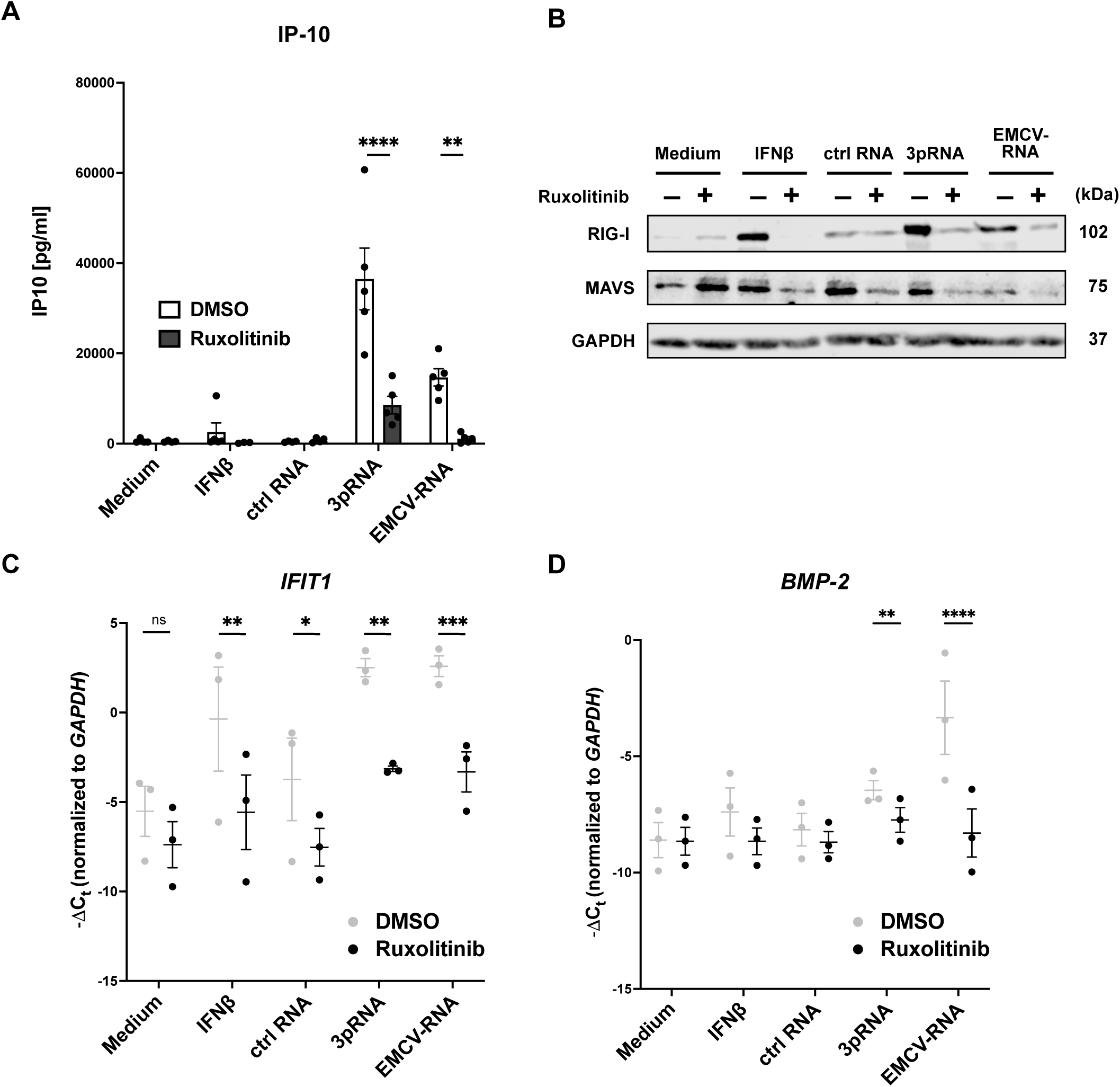
Inhibition of JAK-STAT pathway by ruxolitinib inhibits RIG-I-like receptor signaling including the induction of osteogenic BMP-2 in human aortic smooth muscle cells. Human aortic smooth muscle cells (HAoSMCs) were preincubated with the JAK-STAT inhibitor ruxolitinib or DMSO (control) for 45 min and subsequently stimulated with IFNβ, ctrl RNA, 3pRNA or EMCV-RNA. (A) Ruxolitinib inhibited the secretion of RIG-I (3pRNA)- and MDA5 (EMCV-RNA)-induced IP-10 (n=4). (B) Ruxolitinib reduced IFNβ-, RIG-I (3pRNA)- and MDA5 (EMCV-RNA)-induced expression of RIG-I and MAVS protein. Representative western blot membranes are shown. (C) Ruxolitinib inhibited the IFNβ-, RIG-I (3pRNA and MDA5 (EMCV-RNA)-induced mRNA expression of the ISG *IFIT1* (n=3) and (D) the osteogenic factor *BMP-2* (n=3). Data are depicted as individual data points (* p<0.05, ** p<0.01, *** p<0.001, **** p<0.0001, ns = not significant; two-way ANOVA with Sidak’s multiple comparison test; means ± SEM.

In the wire-injury model, C57BL/6J mice were assigned to four groups: sham+chow (n = 5), sham+ruxolitinib (n = 5), AVS+chow (n = 10), AVS+ruxolitinib (n = 10). Mice received ruxolitinib (2 g/kg, ad libitum in chow) or standard chow from three days before surgery through day 42 (Fig. 6A). Echocardiography was performed at days −2, 14, 28, and 42. Ruxolitinib significantly reduced circulating T cells, dendritic cells, and NK cells (percentage of all CD45⁺ cells) compared with chow in the AVS group (Fig. 6B). The peak transvalvular gradient time course showed a significant difference in the temporal trajectories across the four groups (mixed-effects model, time x treatment interaction). The mean gradient showed the same time x treatment interaction (Fig. 6C). At day 42, ruxolitinib-treated AVS mice had a significantly larger systolic area than chow-fed AVS mice, and chow-fed AVS mice in turn had significantly smaller systolic area than chow-fed sham mice, indicating that the injury-related reduction in systolic area was significantly attenuated by ruxolitinib. For the diastolic area, the AVS+chow versus AVS+ruxolitinib comparison at day 42 was also significantly reduced (Fig. 6D). At the earlier day-14 timepoint, significant pairwise differences were confined to the Sham+ruxolitinib subgroup versus the other three arms and did not involve the AVS+chow versus AVS+ruxolitinib comparison, which reached significance at day 42. Thus, ruxolitinib improved the recovery from wire injury rather than reducing the injury-induced AVS induction directly after wire injury. Survival differed significantly across the four groups (log-rank Mantel-Cox test, χ² = 9.05, df = 3, p = 0.029; Fig. 6E). Isolated comparison between the injury-induced AVS groups revealed significantly higher survival of the AVS+ruxolitinib mice compared to the AVS+chow mice (log-rank Mantel-Cox test, χ² = 5.31, df = 1, p = 0.021; Fig. 6E).

**Fig. 6:**
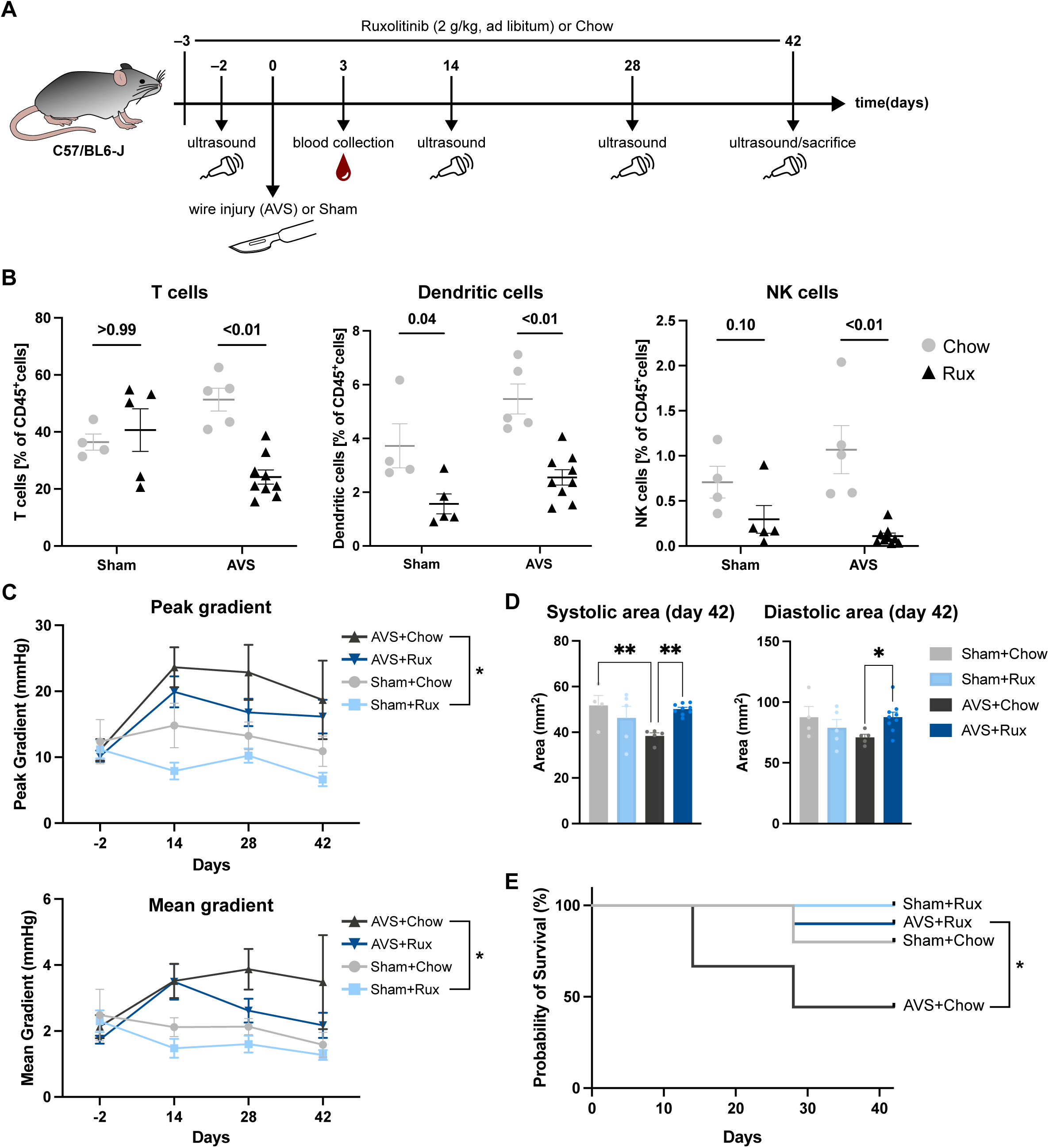
Inhibition of JAK-STAT pathway by ruxolitinib improves recovery from wire-injury-induced aortic valve stenosis. (A) Experimental design of the wire injury model and intervention with the JAK/STAT inhibitor ruxolitinib. C57BL/6J mice were exposed to wire injury (AVS) or sham surgery (Sham) at day 0. Mice received chow food ad libitum either with ruxolitinib at a dose of 2 g/kg chow (Rux) or without ruxolitinib as control (Chow) daily starting three days prior to surgery until mice were sacrificed at day 42 (Sham+Chow n=5, Sham+Rux n=5, AVS+Chow n=10, AVS+Rux n=10). Cardiac ultrasound was performed at the indicated time points. (B) Three days after wire injury, immune cell subsets (T cells, dendritic cells, NK cells) were quantified by flow cytometry. (C) Echocardiography was performed 2 days prior to wire injury and at days 14, 28 and 42 after wire injury. Maximum blood flow velocity (v) across the stenotic outflow region was determined and the maximum pressure gradient calculated (ΔP = 4v²). Higher pressure gradient (peak gradient, upper graph) indicates a higher degree of obstruction and a higher acute pressure load. Higher average pressure gradient during the systolic ejection phase (mean gradient, lower graph) indicates sustained hemodynamic pressure and correlates with a more severe aortic valve stenosis. (D) The systolic (left) and the diastolic (right) area of the left ventricular cavity on day 42. (E) Survival or required removal from experiment due to severity of symptoms. Statistical difference between all groups over time was calculated by using the *mixed-effects model time x treatment* (peak gradient: p < 0.01; mean gradient: p < 0.05). The statistical difference of systolic and diastolic areas on day 42 were determined by *One-way ANOVA with Fisher’s LSD test*. The p-values refer to an overall difference in temporal trajectories across the four groups (* p < 0.05, ** p < 0.01). For survival analysis: two-sided log-rank test; across all groups: p=0.029; between AVS groups (± ruxolitinib): p=0.02; * p < 0.05.

Together, ruxolitinib suppresses RIG-I-dependent type I interferon signaling and osteogenic gene expression *in vitro*. *In vivo*, ruxolitinib reduces circulating innate (NK cells, dendritic cells) and adaptive (T cells) immune cell populations, attenuates hemodynamic disease progression, preserves valve area, and improves survival in the wire-injury model of aortic valve stenosis. These results identify JAK/STAT inhibition as a promising therapeutic strategy for the prevention of disease progression of aortic valve stenosis.

## Discussion

In this study, we show that human aortic interstitial cells are RIG-I-like receptor (RLR)-responsive cells. Activation of RIG-I or MDA5 by exogenous agonists induces a type I interferon response and a G3BP1-dependent osteogenic transcriptional program driving MAVS-dependent calcification *in vitro*. Strikingly, cyclic mechanical stretch alone, in the absence of any exogenous ligand, was found to activate the RIG-I/MAVS-dependent pathway to a certain degree. This activation is caused by the release of mitochondrial double-stranded RNA (dsRNA) which was found to be the dominant endogenous dsRNA species detected. *In vivo*, constitutive RLR signaling contributed to hemodynamic disease progression in a wire-injury model of aortic valve stenosis. Pharmacological JAK/STAT inhibition with ruxolitinib attenuated disease progression, preserved systolic and diastolic area of the left ventricular cavity, and improved survival in this model. These *in vivo* findings were supported by ruxolitinib-mediated suppression of RIG-I-dependent osteogenic signaling *in vitro*. Together, these results identify mechanical-stretch-induced, mitochondrial dsRNA-driven RIG-I activation as a mechanistic link between the biomechanical environment of the aorta and the innate-immune-osteogenic program that drives its calcification, and identify ruxolitinib as a novel promising therapeutic strategy.

These findings resolve the following paradox: In Singleton-Merten syndrome (SMS) constitutive, exogenous ligand-independent RIG-I or MDA5 activation caused by gain of function mutations is sufficient to drive aortic and valvular calcification ^15–18^. However, neither viral infection nor the administration of IFN-α or IFN-β for the treatment of chronic hepatitis B or multiple sclerosis, have been reported to cause aortic or valvular calcification, despite robust systemic type I IFN pathway activation and well-documented cardiovascular toxicity of these regimens ^20–22^. Our data suggest that in SMS the relevant trigger of aortic calcification is not the systemic interferon response itself, but a local, cell-intrinsic and endogenous ligand-dependent activation of RIG-I within the aortic wall. Because in SMS gain of function mutations render RIG-I or MDA5 constitutively active in the absence of ligand, the aortic wall in these patients is presumably constantly exposed to ligand-independent signaling. The data from our work here demonstrate that wild-type RIG-I in aortic wall cells can be activated locally through mechanical stretch of aortic interstitial cells leading to leakage of endogenous mitochondrial dsRNA into the cytoplasm. This raises the possibility that any process capable of triggering local RIG-I activation in the aortic wall, including but not limited to germline gain-of-function mutations, converges on the same osteogenic output, whereas systemic activation of a type I IFN response without a local aortic wall trigger does not. Notably, recognition of the endogenous ligand mitochondrial dsRNA was RIG-I dependent, showing that under physiological mechanical stress conditions in the aortic wall, constitutive RIG-I activation contributes to osteogenic signaling.

The precise molecular mechanism connecting mechanical stress to RIG-I activation remains incompletely defined by the present data and warrants further investigation. We show that mitochondrial RNA is the dominant dsRNA species detectable in mechanically stressed aortic interstitial cells, and that its pharmacological depletion with the mitochondrial RNA polymerase inhibitor IMT1 was associated with loss of the stretch-induced ISG response. However, it is not yet established at what point in the mechanotransduction cascade mitochondrial dsRNA is released into the cytosol, nor precisely where within the cell RIG-I engages this ligand. Furthermore, the exact ligand structure itself is not defined yet. It is well-established that the RIG-I binding pocket recognizes 5′-triphosphate or 5′-diphosphate blunt-ended dsRNA^4,5^. However, the great majority of mitochondrial dsRNA in fact carries a 5′-monophosphate rather than a canonical 5′-triphosphate terminus and is therefore excluded from recognition by RIG-I ^10^. How mechanical stress might generate and expose RIG-I-ligand competent RNA from the mitochondrial RNA pool remains unclear and represents an important direction for follow-up work. Our data are consistent with previous studies that reported RIG-I-dependent recognition of mt-dsRNA leaking into the cytosol ^9,12–14^. Leakage of mtRNA has been found to be triggered by various mechanisms including impairment of the mitochondrial RNA degradasome ^9,13^, modulation of the tricarboxylic acid cycle ^14,49^ and disruption of mitochondrial dynamics ^12^ in a mitochondrial-pore (BAX/BAK or VDAC1) dependent way. Moreover, mechanical stress like cyclic stretch has been described to directly affect mitochondrial dynamics (e.g., morphology, fission and fusion events; reviewed in ^50^. Therefore, impairment of mitochondrial dynamics by mechanical stress is a likely candidate for the mechanism of mt-dsRNA leakage to the cytosol and consequent RIG-I activation.

We also considered the phosphatase PP1 which has been implicated in cytoskeletal-stress-associated RLR priming through the PP1 regulatory subunit PPP1R12C ^48^. PP1 has been reported to dephosphorylate and thereby license RIG-I for ligand-induced activation ^47^, thereby potentially providing a mechanistic bridge between mechanical stress, which directly affects the cytoskeletal architecture ^51^ and RIG-I activation. We were not able to experimentally reproduce a role for PP1 in mechanical-stress-induced RIG-I co-activation in aortic interstitial cells (unpublished results). Nevertheless, a PP1-dependent, cytoskeleton-linked priming mechanism remains an attractive hypothesis for how mechanical force is transduced into RLR pathway engagement, and future work with different experimental approaches may provide evidence for PP1 involvement under certain environmental and cellular conditions.

Downstream of RIG-I/MAVS, we identified the stress-granule protein G3BP1 as a requirement for RIG-I-driven osteogenic gene expression and calcification. This is consistent with a growing body of evidence that G3BP1 functions as a signaling node connecting nucleic-acid sensing to downstream transcriptional and translational programs. G3BP1 is required for efficient translation of interferon-stimulated transcripts during stress-granule-associated translational reprogramming ^35^, and it promotes cGAS-dependent DNA sensing ^36^. Furthermore, G3BP1 has been shown to convey RIG-I/MAVS-dependent, pro-sclerotic non-canonical Wnt signals in SMS-mutant vascular smooth muscle cells by cooperating with Dishevelled independently of canonical β-catenin signaling ^37,38^. Our finding that FZD5, a Wnt receptor previously implicated in Wnt5a-driven vascular calcification ^52,53^, is likewise required for RLR-driven calcification is consistent with this non-canonical Wnt axis operating downstream of G3BP1 in HAoSMCs, although we did not formally test whether FZD5 acts within the same G3BP1-dependent signaling module or in a parallel pathway. Together with the ruxolitinib data discussed below, these results position G3BP1, and potentially FZD5, as candidate therapeutic targets acting downstream of RIG-I activation, with the practical advantage that intervening downstream of RIG-I could in principle limit osteogenic signaling without broadly suppressing its antiviral function.

Our results extend a body of literature implicating nucleic-acid-sensing pattern-recognition receptors, including but not limited to RLRs, in vascular and valvular calcification. Toll-like receptors have previously been shown to contribute to calcific aortic valve disease and atherosclerosis: TLR3 activation induces osteogenic responses in human aortic valve interstitial cells ^30^ and contributes to aortic valve stenosis *in vivo* ^27^, TLR3/TLR4 signaling through TRIF/TRAM promotes atherosclerosis ^54^, and RIG-I itself has been detected in atherosclerotic macrophages, with the receptor proposed as a contributor to atherogenesis on largely correlative grounds ^28^. Our data build on this literature by providing a direct, genetically and pharmacologically supported causal link between RIG-I/MAVS signaling and osteogenic transdifferentiation and calcification specifically in aortic smooth muscle cells. To our knowledge, this is the first study reporting that this osteogenic pathway can be triggered by mechanical stress in the absence of any exogenous or microbial ligand. This distinguishes our findings from prior work implicating RLRs or TLRs in calcification predominantly in the context of infection, sterile DNA damage, or lipid-driven inflammation ^29,55–57^. Instead, our findings place biomechanical load itself upstream of innate immune activation in the aortic valve.

Our *in vivo* data support a functional contribution of this pathway to disease in a physiologically relevant model. MAVS deficiency significantly attenuated hemodynamic disease progression after wire injury, and pharmacological JAK/STAT inhibition with ruxolitinib reduced circulating innate and adaptive immune cell populations, attenuated hemodynamic disease progression, preserved the area of the left ventricular cavity, and improved survival. Ruxolitinib is an already-approved, orally available JAK1/2 inhibitor with an established long-term safety profile in myeloproliferative disease. In pediatric interferonopathies ruxolitinib has shown efficacy in conditions related to SMS and in STING-associated vasculopathy of infancy ^39,40^. Since a related JAK inhibitor has independently been shown to blunt dsRNA-induced inflammation and calcification in human aortic valve interstitial cells ^58^, our data support clinical testing of ruxolitinib as a repurposable candidate therapeutic for aortic valve stenosis. Of note, JAK inhibition may also affect osteogenic differentiation through RIG-I-independent mechanisms, as ruxolitinib has been shown to influence osteoblastic differentiation of human skeletal stem cells in an unrelated screening context ^59^. This should be considered when interpreting the beneficial effect in a future clinical trial, which may reflect both RIG-I-pathway-specific as well as broader anti-osteogenic actions of the drug.

Several limitations should be considered when interpreting the findings of our study. First, our functional *in vivo* data rely on one wire-injury cohort and, for the ruxolitinib intervention, on group sizes of 5 to 10 animals per arm. Second, and more broadly, RLR receptor biology shows species differences, and it remains uncertain whether such discrepancies reflect genuinely redundant sensing mechanisms or true differences between mouse and human RLR biology ^60^. Because our *in vivo* functional data were generated in mice while our mechanistic and cell-biological data were generated in primary human cells, the species difference is a relevant caveat when extrapolating the combined dataset to human aortic valve stenosis.

In conclusion, we show that mechanical-stress-induced, mitochondrial-dsRNA-associated activation of RIG-I drives a G3BP1-dependent osteogenic program in aortic smooth muscle cells that presumably contributes to aortic valve stenosis *in vivo*, and that this pathway is pharmacologically tractable with the JAK/STAT inhibitor ruxolitinib. These findings are consistent with, and provide a plausible physiological counterpart to, the genetic proof of concept provided by Singleton-Merten syndrome, in which constitutive RIG-I or MDA5 activation alone is sufficient to cause aortic and valvular calcification. At the same time, our data do not establish the quantitative contribution of this RIG-I-dependent mechanism relative to other, non-RLR-dependent pathogenic and metabolic drivers of aortic valve calcification, such as hyperphosphatemia-driven mineral nucleation, BMP-2/Runx2-driven osteogenic transdifferentiation acting independently of innate immune signaling, or lipid- and TLR-driven inflammatory calcification ^23,25,60^. Whether environmental triggers such as subclinical infection, additional genetic modifiers of RIG-I pathway activity, or metabolic cofactors act together with mechanical load to determine the extent of RIG-I-dependent calcification in an individual aortic valve remains an open question that will require further study.

## Acknowledgements

This study was funded by the Deutsche Forschungsgemeinschaft (DFG, German Research Foundation) under Germany’s Excellence Strategy – EXC2151 – 390873048 of which KT, SS, SZ, RPN, GN, EB and GH are members. It was also supported by other grants of the DFG, including TRR259 (EB, GH, SS, PRG, SZ and GN; project-ID 397484323); TRR237 (EB, GH, MS, TZ; project-ID 369799452). This work is part of the PhD theses of Madeleine Gräf and Paraskevi Vasileiadou at the University of Bonn. The authors gratefully acknowledge the granted access to the Marvin and Bonna clusters hosted by the University of Bonn. We would like to acknowledge the central facility S01 Animal Models of the TRR259, as well as the assistance of the Next Generation Sequencing and Histology Core Facilities of the Medical Faculty at the University of Bonn.

MG, PV, FB and SoS performed the experiments. SS, MS, TZ, EB and GH conceived experiments. SD, FB, TSG, LS, JS, KM, KT, RPN, SoS, PRG, SZ, GN, EB and GH analyzed and interpreted data. MG, EB and GH wrote the manuscript. All authors revised the manuscript.

## Data Availability

The data that support the findings of this study are available from the corresponding author upon reasonable request.

## References

1. Yoneyama M, Kikuchi M, Natsukawa T, Shinobu N, Imaizumi T, Miyagishi M, Taira K, Akira S, Fujita T. The RNA helicase RIG-I has an essential function in double-stranded RNA-induced innate antiviral responses. Nat. Immunol. 2004;5:730–737.

2. Kato H, Takeuchi O, Sato S, Yoneyama M, Yamamoto M, Matsui K, Uematsu S, Jung A, Kawai T, Ishii KJ, et al. Differential roles of MDA5 and RIG-I helicases in the recognition of RNA viruses. Nature. 2006;441:101–105.

3. Yoneyama M, Kikuchi M, Matsumoto K, Imaizumi T, Miyagishi M, Taira K, Foy E, Loo Y-M, Gale M, Akira S, et al. Shared and Unique Functions of the DExD/H-Box Helicases RIG-I, MDA5, and LGP2 in Antiviral Innate Immunity. J. Immunol. 2005;175:2851–2858.

4. Hornung V, Ellegast J, Kim S, Brzózka K, Jung A, Kato H, Poeck H, Akira S, Conzelmann K-K, Schlee M, et al. 5’-Triphosphate RNA Is the Ligand for RIG-I. Science. 2006;314:994–997.

5. Goubau D, Schlee M, Deddouche S, Pruijssers AJ, Zillinger T, Goldeck M, Schuberth C, Veen AGV der, Fujimura T, Rehwinkel J, et al. Antiviral immunity via RIG-I-mediated recognition of RNA bearing 5′-diphosphates. Nature. 2014;514:372–375.

6. Schlee M, Hartmann E, Coch C, Wimmenauer V, Janke M, Barchet W, Hartmann G. Approaching the RNA ligand for RIG-I? Immunol. Rev. 2009;227:66–74.

7. Schlee M, Roth A, Hornung V, Hagmann CA, Wimmenauer V, Barchet W, Coch C, Janke M, Mihailovic A, Wardle G, et al. Recognition of 5’ triphosphate by RIG-I helicase requires short blunt double-stranded RNA as contained in panhandle of negative-strand virus. Immunity. 2009;31:25–34.

8. Young PG, Attardi G. Characterization of double-stranded RNA from HeLa cell mitochondria. Biochem. Biophys. Res. Commun. 1975;65:1201–1207.

9. Dhir A, Dhir S, Borowski LS, Jimenez L, Teitell M, Rötig A, Crow YJ, Rice GI, Duffy D, Tamby C, et al. Mitochondrial double-stranded RNA triggers antiviral signalling in humans. Nature. 2018;560:238–242.

10. Regt AK de, Anand K, Ciupka K, Bender F, Gatterdam K, Putschli B, Fusshöller D, Hilbig D, Kirchhoff A, Hunkler C, et al. A conserved isoleucine in the binding pocket of RIG-I controls immune tolerance to mitochondrial RNA. Nucleic acids Res. 2023;51:11893–11910.

11. Regt AK de, Anand K, Ciupka K, Bender F, Gatterdam K, Putschli B, Fusshöller D, Hilbig D, Kirchhoff A, Hunkler C, et al. A conserved isoleucine in the binding pocket of RIG-I controls immune tolerance to mitochondrial RNA. Nucleic acids Res. 2023;51:11893–11910.

12. Yasuda T, Ichikawa A, Onoue K, Ogasawara E, Ishihara T, Kosako H, Ishihara N. Disrupted mitochondrial dynamics activate RNA-sensing innate immunity through mitochondrial RNA release. Cell Rep. 2026;45:117607.

13. López-Polo V, Maus M, Zacharioudakis E, Lafarga M, Attolini CS-O, Marques FDM, Kovatcheva M, Gavathiotis E, Serrano M. Release of mitochondrial dsRNA into the cytosol is a key driver of the inflammatory phenotype of senescent cells. Nat. Commun. 2024;15:7378.

14. Hooftman A, Peace CG, Ryan DG, Day EA, Yang M, McGettrick AF, Yin M, Montano EN, Huo L, Toller-Kawahisa JE, et al. Macrophage fumarate hydratase restrains mtRNA-mediated interferon production. Nature. 2023;615:490–498.

15. Jang M-A, Kim EK, Now H, Nguyen NTH, Kim W-J, Yoo J-Y, Lee J, Jeong Y-M, Kim C-H, Kim O-H, et al. Mutations in DDX58, which Encodes RIG-I, Cause Atypical Singleton-Merten Syndrome. Am. J. Hum. Genet. 2015;96:266–274.

16. Rutsch F, MacDougall M, Lu C, Buers I, Mamaeva O, Nitschke Y, Rice GI, Erlandsen H, Kehl HG, Thiele H, et al. A Specific IFIH1 Gain-of-Function Mutation Causes Singleton-Merten Syndrome. Am. J. Hum. Genet. 2015;96:275–282.

17. Lässig C, Lammens K, López JLG, Michalski S, Fettscher O, Hopfner K-P. Unified mechanisms for self-RNA recognition by RIG-I Singleton-Merten syndrome variants. eLife. 2018;7:e38958.

18. Singleton EB, Merten DF. An unusual syndrome of widened medullary cavities of the metacarpals and phalanges, aortic calcification and abnormal dentition. Pediatr. Radiol. 1973;1:2–7.

19. Lu C, MacDougall M. RIG-I-Like Receptor Signaling in Singleton-Merten Syndrome. Front. Genet. 2017;8:118.

20. Reder AT, Ebers GC, Traboulsee A, Li D, Langdon D, Goodin DS, Bogumil T, Beckmann K, Konieczny A, Study I of the 16-YL-TF-U. Cross-sectional study assessing long-term safety of interferon-&bgr;-1b for relapsing-remitting MS(LOE Classification). Neurology. 2010;74:1877–1885.

21. Gold R, Rieckmann P, Chang P, Abdalla J, Group the PS. The long-term safety and tolerability of high-dose interferon β-1a in relapsing–remitting multiple sclerosis: 4-year data from the PRISMS study. Eur. J. Neurol. 2005;12:649–656.

22. Genentech Inc. Pegasys (peginterferon alfa-2a) prescribing information [Internet]. South San Francisco, CA: 2023.

23. Durham AL, Speer MY, Scatena M, Giachelli CM, Shanahan CM. Role of smooth muscle cells in vascular calcification: implications in atherosclerosis and arterial stiffness. Cardiovasc. Res. 2018;114:590–600.

24. Cao G, Xuan X, Hu J, Zhang R, Jin H, Dong H. How vascular smooth muscle cell phenotype switching contributes to vascular disease. Cell Commun. Signal. 2022;20:180.

25. Boström K, Watson KE, Horn S, Wortham C, Herman IM, Demer LL. Bone morphogenetic protein expression in human atherosclerotic lesions. J. Clin. Investig. 1993;91:1800–1809.

26. Ducy P, Zhang R, Geoffroy V, Ridall AL, Karsenty G. Osf2/Cbfa1: A Transcriptional Activator of Osteoblast Differentiation. Cell. 1997;89:747–754.

27. Niepmann ST, Willemsen N, Boucher AS, Stei M, Goody P, Zietzer A, Bulic M, Billig H, Odainic A, Weisheit CK, et al. Toll-like receptor-3 contributes to the development of aortic valve stenosis. Basic Res. Cardiol. 2023;118:6.

28. Imaizumi T, Yagihashi N, Kubota K, Yoshida H, Sakaki H, Yagihashi S, Kimura H, Satoh K. Expression of Retinoic Acid-inducible gene-I (RIG-I) in Macrophages: Possible Involvement of RIG-I in Atherosclerosis. J. Atheroscler. Thromb. 2007;14:51.

29. Ahmad U, Ali R, Lebastchi AH, Qin L, Lo SL, Yakimov AO, Khan SF, Choy JC, Geirsson A, Pober JS, et al. IFN-γ Primes Intact Human Coronary Arteries and Cultured Coronary Smooth Muscle Cells to Double-Stranded RNA- and Self-RNA–Induced Inflammatory Responses by Upregulating TLR3 and Melanoma Differentiation-Associated Gene 5. J. Immunol. 2010;185:1283–1294.

30. Zhan Q, Song R, Zeng Q, Yao Q, Ao L, Xu D, Fullerton DA, Meng X. Activation of TLR3 Induces Osteogenic Responses in Human Aortic Valve Interstitial Cells through the NF-κB and ERK1/2 Pathways. Int. J. Biol. Sci. 2015;11:482–493.

31. Bäck M, Gasser TC, Michel J-B, Caligiuri G. Biomechanical factors in the biology of aortic wall and aortic valve diseases. Cardiovasc. Res. 2013;99:232–241.

32. Jensen LF, Bentzon JF, Albarrán-Juárez J. The Phenotypic Responses of Vascular Smooth Muscle Cells Exposed to Mechanical Cues. Cells. 2021;10:2209.

33. Isnard RN, Pannier BM, Laurent S, London GM, Diebold B, Safar ME. Pulsatile diameter and elastic modulus of the aortic arch in essential hypertension: A noninvasive study. J. Am. Coll. Cardiol. 1989;13:399–405.

34. Anwar MA, Shalhoub J, Lim CS, Gohel MS, Davies AH. The Effect of Pressure-Induced Mechanical Stretch on Vascular Wall Differential Gene Expression. J. Vasc. Res. 2012;49:463–478.

35. Bidet K, Dadlani D, Garcia-Blanco MA. G3BP1, G3BP2 and CAPRIN1 Are Required for Translation of Interferon Stimulated mRNAs and Are Targeted by a Dengue Virus Non-coding RNA. PLoS Pathog. 2014;10:e1004242.

36. Liu Z-S, Cai H, Xue W, Wang M, Xia T, Li W-J, Xing J-Q, Zhao M, Huang Y-J, Chen S, et al. G3BP1 promotes DNA binding and activation of cGAS. Nat. Immunol. 2019;20:18–28.

37. Ramachandran B, Stabley JN, Cheng S-L, Behrmann AS, Gay A, Li L, Mead M, Kozlitina J, Lemoff A, Mirzaei H, et al. A GTPase-activating protein–binding protein (G3BP1)/antiviral protein relay conveys arteriosclerotic Wnt signals in aortic smooth muscle cells. J. Biol. Chem. 2018;293:7942–7968.

38. Bikkavilli RK, Malbon CC. Arginine methylation of G3BP1 in response to Wnt3a regulates β-catenin mRNA. J. Cell Sci. 2011;124:2310–2320.

39. Frémond M-L, Rodero MP, Jeremiah N, Belot A, Jeziorski E, Duffy D, Bessis D, Cros G, Rice GI, Charbit B, et al. Efficacy of the Janus kinase 1/2 inhibitor ruxolitinib in the treatment of vasculopathy associated with TMEM173-activating mutations in 3 children. J. Allergy Clin. Immunol. 2016;138:1752–1755.

40. Broser P, Mengershausen U von, Heldt K, Bartholdi D, Braun D, Wolf C, Lee-Kirsch MA. Precision treatment of Singleton Merten syndrome with ruxolitinib: a case report. Pediatr. Rheumatol. 2022;20:24.

41. Niepmann ST, Steffen E, Zietzer A, Adam M, Nordsiek J, Gyamfi-Poku I, Piayda K, Sinning J-M, Baldus S, Kelm M, et al. Graded murine wire-induced aortic valve stenosis model mimics human functional and morphological disease phenotype. Clin. Res. Cardiol. 2019;108:847–856.

42. Goldeck M, Schlee M, Hartmann G, Hornung V. Innate DNA and RNA Recognition, Methods and Protocols. Methods Mol. Biol. 2014;1169:15–25.

43. Brinkman EK, Chen T, Amendola M, Steensel B van. Easy quantitative assessment of genome editing by sequence trace decomposition. Nucleic Acids Res. 2014;42:e168–e168.

44. Love MI, Huber W, Anders S. Moderated estimation of fold change and dispersion for RNA-seq data with DESeq2. Genome Biol. 2014;15:550.

45. Stephens M. False discovery rates: a new deal. Biostatistics. 2017;18:275–294.

46. Wu T, Hu E, Xu S, Chen M, Guo P, Dai Z, Feng T, Zhou L, Tang W, Zhan L, et al. clusterProfiler 4.0: A universal enrichment tool for interpreting omics data. Innov. 2021;2:100141.

47. Wickham H, Averick M, Bryan J, Chang W, McGowan L, François R, Grolemund G, Hayes A, Henry L, Hester J, et al. Welcome to the Tidyverse. J. Open Source Softw. 2019;4:1686.

48. Bender F, Dudek S, Hega M, Maser K, Awaragi A, Putschli B, Wieseler J, Behrendt R, Kümmerer B, Hartmann G, et al. SALDOR-Seq: a novel workflow for full-length sequencing of long dsRNA. ResearchSquare. 2026;

49. O’Carroll SM, Peace CG, Toller-Kawahisa JE, Min Y, Hooftman A, Charki S, Kehoe L, O’Sullivan MJ, Zoller A, Mcgettrick AF, et al. Itaconate drives mtRNA-mediated type I interferon production through inhibition of succinate dehydrogenase. Nat. Metab. 2024;6:2060–2069.

50. Prakash A, Iskratsch T. Mitochondrial Adaptation to Mechanical Stress in Cardiac Ageing and Disease. Adv. Sci. 2026;13:e16736.

51. Fletcher DA, Mullins RD. Cell mechanics and the cytoskeleton. Nature. 2010;463:485–492.

52. Woldt E, Terrand J, Mlih M, Matz RL, Bruban V, Coudane F, Foppolo S, Asmar ZE, Chollet ME, Ninio E, et al. The nuclear hormone receptor PPARγ counteracts vascular calcification by inhibiting Wnt5a signalling in vascular smooth muscle cells. Nat. Commun. 2012;3:1077.

53. XIN H, XIN F, ZHOU S, GUAN S. The Wnt5a/Ror2 pathway is associated with determination of the differentiation fate of bone marrow mesenchymal stem cells in vascular calcification. Int. J. Mol. Med. 2013;31:583–588.

54. Lundberg AM, Ketelhuth DFJ, Johansson ME, Gerdes N, Liu S, Yamamoto M, Akira S, Hansson GK. Toll-like receptor 3 and 4 signalling through the TRIF and TRAM adaptors in haematopoietic cells promotes atherosclerosis. Cardiovasc. Res. 2013;99:364–373.

55. Zimmer S, Steinmetz M, Asdonk T, Motz I, Coch C, Hartmann E, Barchet W, Wassmann S, Hartmann G, Nickenig G. Activation of Endothelial Toll-Like Receptor 3 Impairs Endothelial Function. Circ. Res. 2011;108:1358–1366.

56. Karadimou G, Plunde O, Pawelzik S-C, Carracedo M, Eriksson P, Franco-Cereceda A, Paulsson-Berne G, Bäck M. TLR7 Expression Is Associated with M2 Macrophage Subset in Calcific Aortic Valve Stenosis. Cells. 2020;9:1710.

57. Luo W, Wang Y, Zhang L, Ren P, Zhang C, Li Y, Azares AR, Zhang M, Guo J, Ghaghada KB, et al. Critical role of cytosolic DNA and its sensing adaptor STING in aortic degeneration, dissection, and rupture. Circulation. 2020;141:42–66.

58. Parra-Izquierdo I, Sánchez-Bayuela T, Castaños-Mollor I, López J, Gómez C, Román JAS, Crespo MS, García-Rodríguez C. Clinically used JAK inhibitor blunts dsRNA-induced inflammation and calcification in aortic valve interstitial cells. FEBS J. 2021;288:6528–6542.

59. AlMuraikhi N, Ali D, Alshanwani A, Vishnubalaji R, Manikandan M, Atteya M, Siyal A, Alfayez M, Aldahmash A, Kassem M, et al. Stem cell library screen identified ruxolitinib as regulator of osteoblastic differentiation of human skeletal stem cells. Stem Cell Res. Ther. 2018;9:319.

60. Choi MK, Wang Z, Ban T, Yanai H, Lu Y, Koshiba R, Nakaima Y, Hangai S, Savitsky D, Nakasato M, et al. A selective contribution of the RIG-I-like receptor pathway to type I interferon responses activated by cytosolic DNA. Proc. Natl. Acad. Sci. 2009;106:17870–17875.

