## Supplementary material for "Mechanical stress-induced release of mitochondrial double-stranded RNA activates RIG-I to drive osteogenic differentiation in aortic disease": SupplFig1

**A**

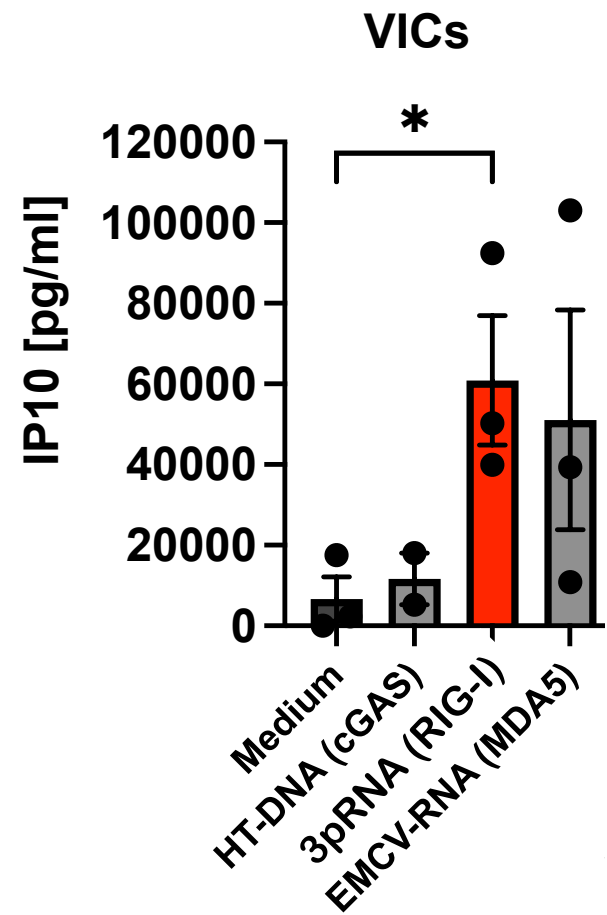

**B**

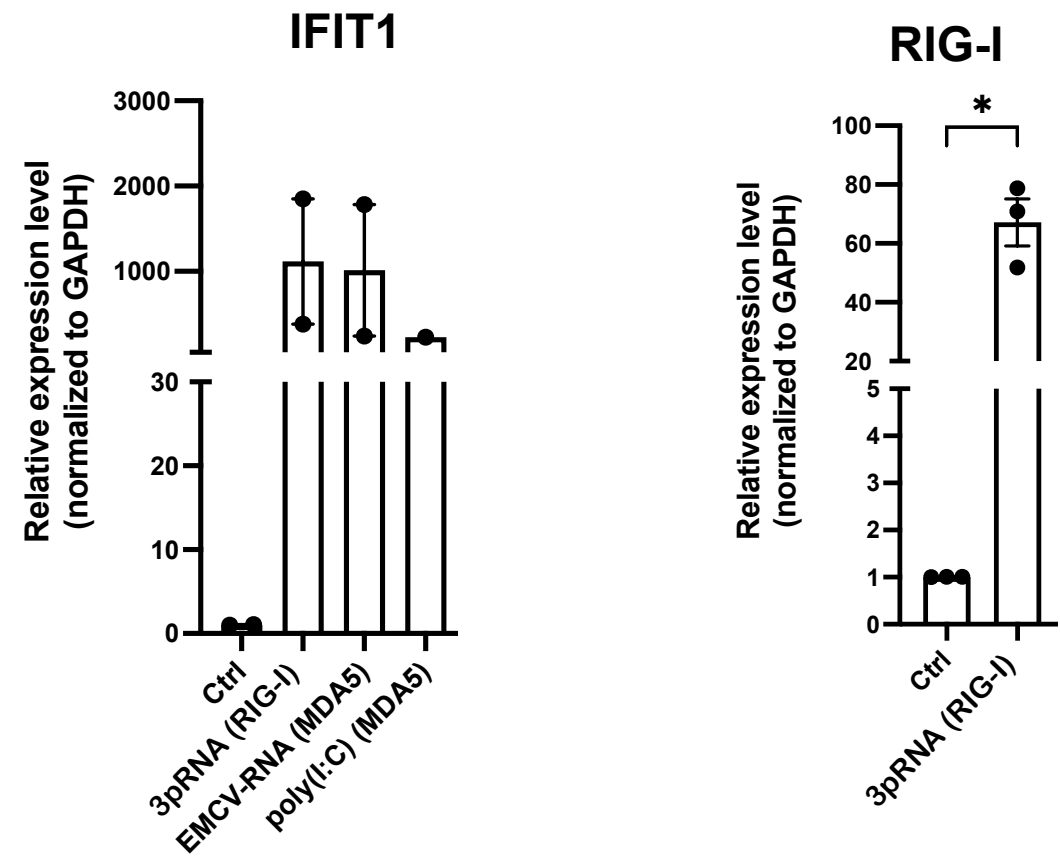

**Supplementary Figure 1: RIG-I activation of human valve interstitial cells (VIC) induces the expression of interferon-stimulated genes including RIG-I.** Human aortic valve tissue was obtained from patients undergoing aortic valve replacement. VICs were prepared and cultured in VIC medium on collagen-coated flasks. VICs were stimulated with 3pRNA (IVT4, RIG-I agonist), EMCV-RNA (MDA5 agonist), poly(I:C) (MDA5 agonist) or HT-DNA (hering testis DNA, cGAS agonist) as indicated. (A) After 24 h, IP-10 was measured in the supernatants by ELISA (n=3 donors). (B) Expression of the interferon-stimulated genes *IFIT1* and *DDX58* (RIG-I) was analysed by qRT-PCR (n=2 donors for *IFIT1* and n=3 donors for *RIG-I*). Data are shown as mean ± SEM. Statistical significance was calculated by one-way ANOVA (A) or Friedman test (B). \* p<0.05.
