## Supplementary material for "Mechanical stress-induced release of mitochondrial double-stranded RNA activates RIG-I to drive osteogenic differentiation in aortic disease": SupplFig2

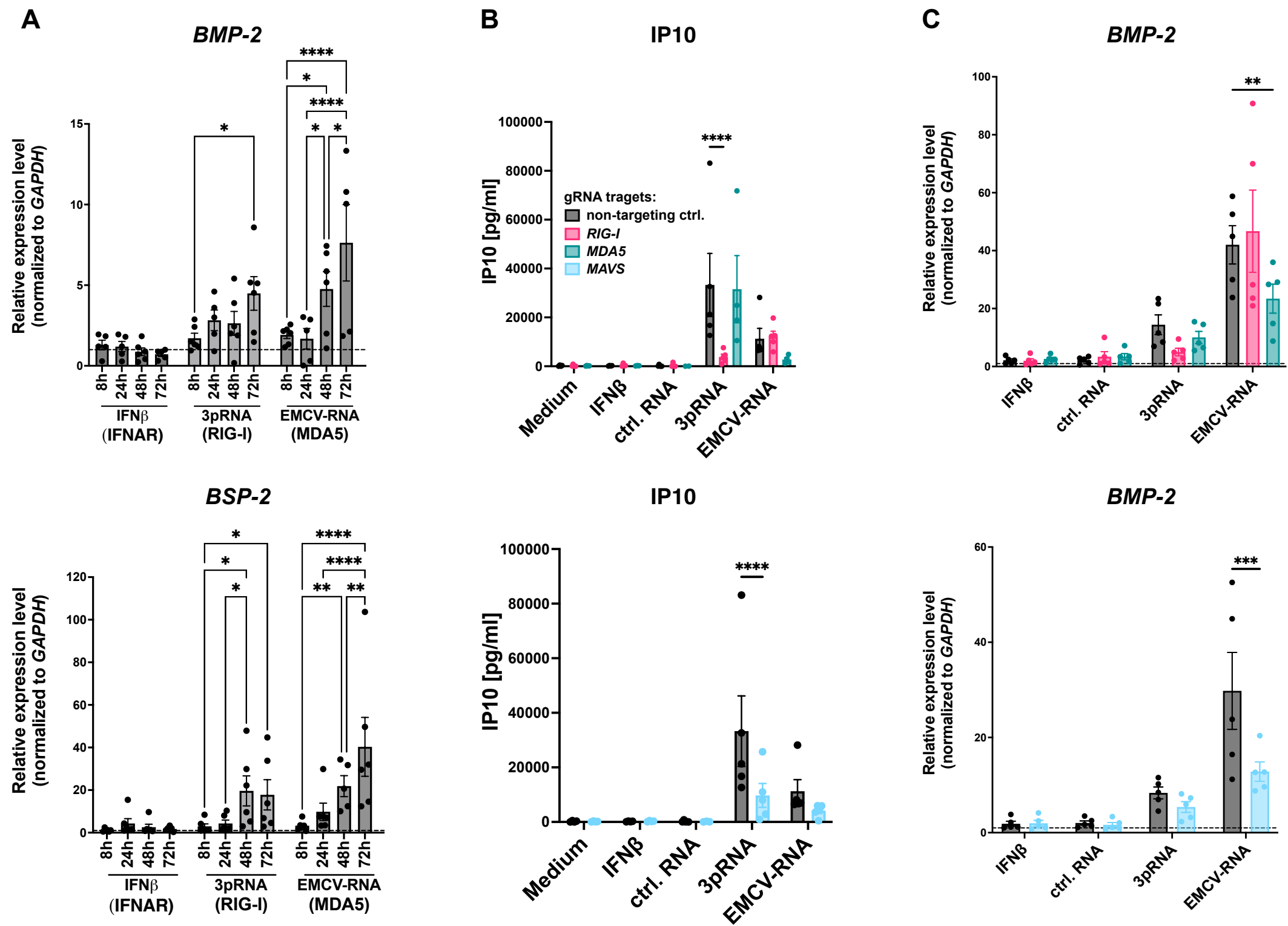

**Supplementary Figure 2. RIG-I-like receptor-dependent upregulation of osteogenic factors in human aortic smooth muscle cells.** HAoSMC were incubated in the presence of RIG-I agonist (3pRNA), MDA5 agonist (EMCV-RNA), control RNA (ctrl RNA), IFNβ or medium. (A) After 72 hours, expression of the osteogenic factors BMP-2 and BSP2 was analyzed by qRT-PCR (n=6 donors). (B) RIG-I deficient HAoSMCs (RIG-I-specific gRNA) and MDA5-deficient HAoSMCs (MDA5-specific gRNA) (upper panel) or MAVS-deficient HAoSMCs (MAVS-specific gRNA) (lower panel) and control HAoSMCs (non-targeting control gRNA) were stimulated for 72 hours, and IP-10 was measured in the supernatants by ELISA (n=4-5 donors for RIG-I- and MDA5-deficient cells; n=5 donors for MAVS-deficient cells). (C) Conditions and cells as in (B), but mRNA expression of BMP-2 was analyzed by qRT-PCR (n=5 donors). Data are shown as mean ± SEM. \* p<0.05, \*\* p<0.01, \*\*\* p<0.001, \*\*\*\* p<0.0001, ns = not significant; statistical significance was calculated by two-way ANOVA with Tukey's multiple comparison test (A) or two-way ANOVA with Dunnett's multiple comparison test (B, C).
