## Supplementary material for "Mechanical stress-induced release of mitochondrial double-stranded RNA activates RIG-I to drive osteogenic differentiation in aortic disease": SupplFig3

**A**

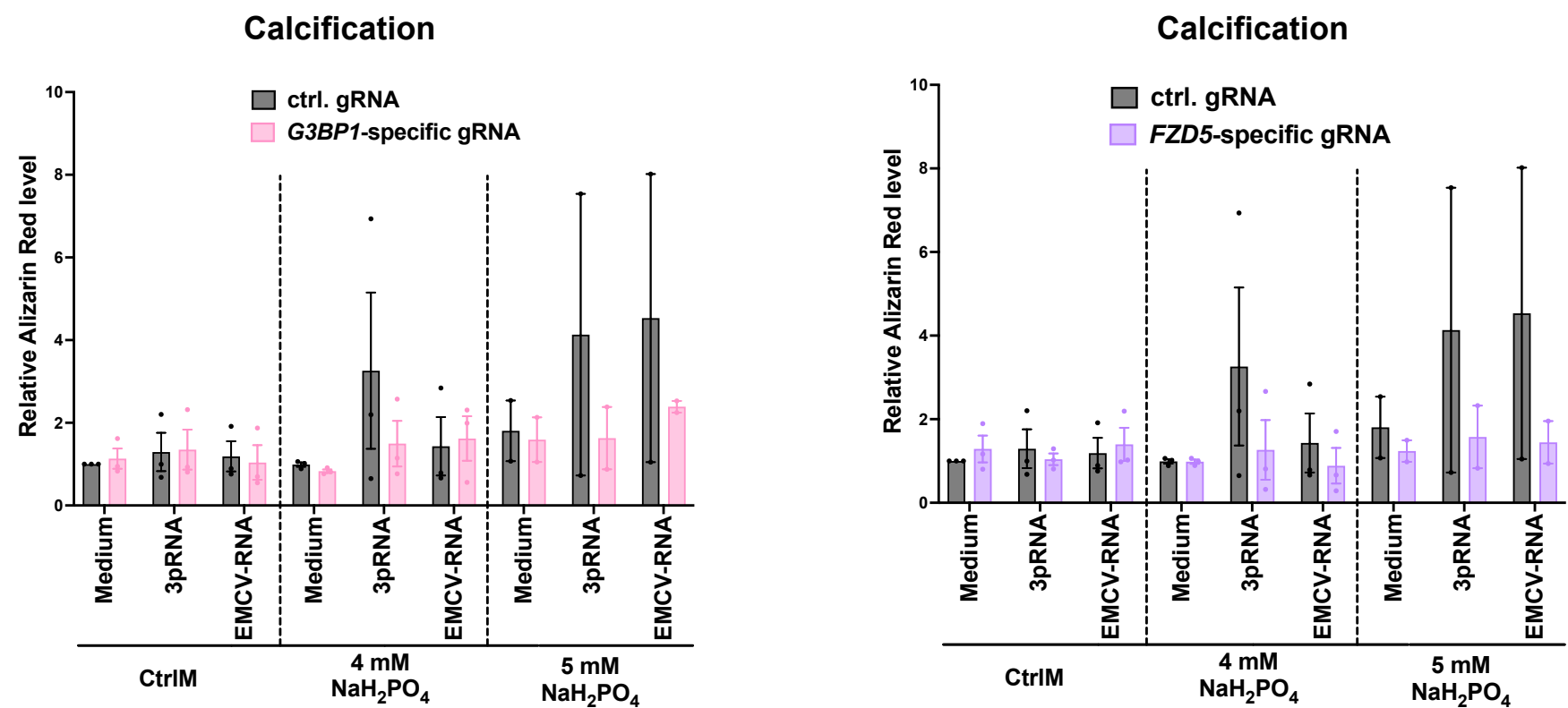

**B**

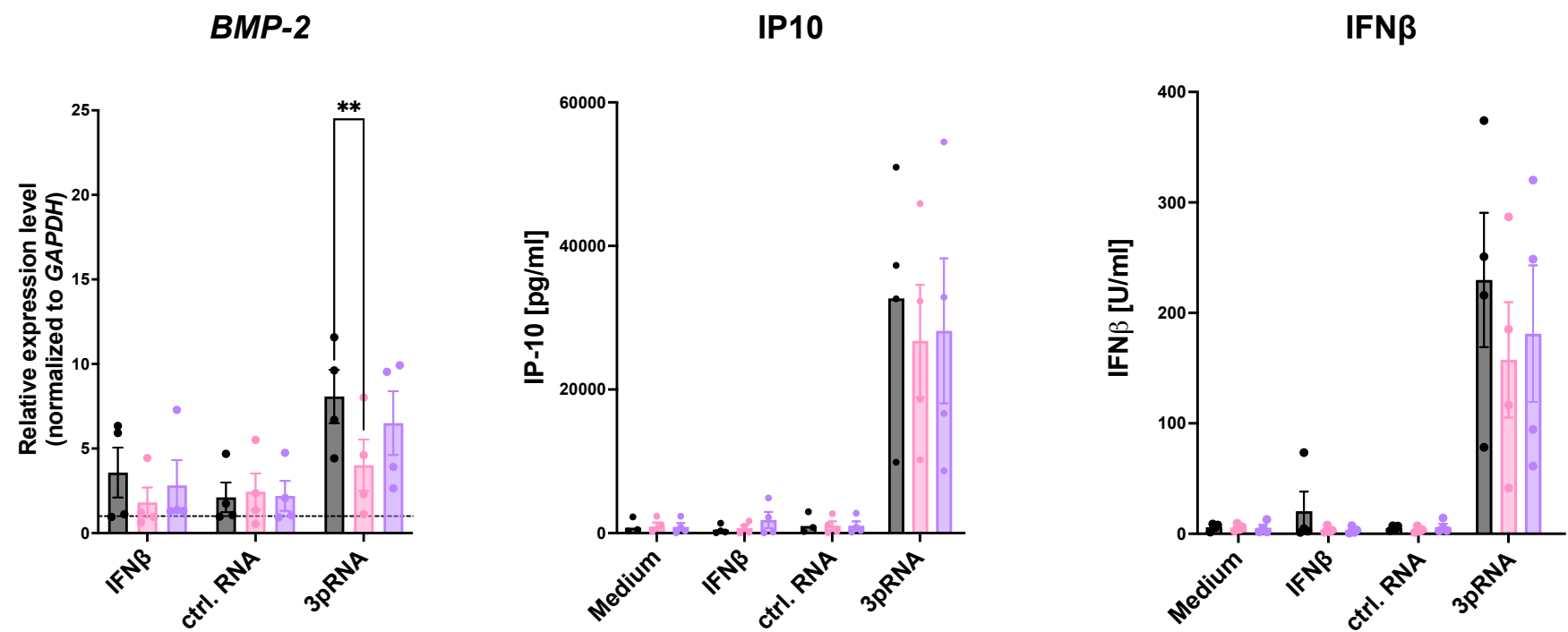

**Supplementary Figure 3: G3BP1 is required for RIG-I-induced upregulation of BMP-2 and calcification.** (A) G3BP1-deficient HAoSMCs (*G3BP1*-specific gRNA), FZD5-deficient HAoSMCs (*FZD5*-specific gRNA) or control HAoSMCs (non-targeting control gRNA) were stimulated with 3pRNA (RIG-I), EMCV-RNA (MDA5), or medium (control) and then cultured in pro-calcifying medium containing 50 μg ascorbic acid and 4 or 5 mM NaH<sub>2</sub>PO<sub>4</sub>, or in control medium. After 7 days, calcium deposition was quantified by Alizarin Red staining. Alizarin Red was dissolved in hexadecylpyridinium chloride (CPC) and the absorbance was measured at 550 nm. (B) G3BP1 deficient HAoSMCs (*G3BP1*-specific gRNA), FZD5-deficient HAoSMCs (*FZD5*-specific gRNA) or control HAoSMCs (non-targeting control gRNA) were stimulated with RIG-I ligand (3pRNA), IFNβ or control RNA. After 72 hours, *BMP-2* mRNA, IP-10 protein and IFNβ protein were quantified (n=4). Data are shown as mean ± SEM. \*\* p<0.01, ns = not significant; statistical significance was calculated by two-way ANOVA with Dunnett's multiple comparison test (B).
