## Supplementary material for "Mechanical stress-induced release of mitochondrial double-stranded RNA activates RIG-I to drive osteogenic differentiation in aortic disease": SupplFig4

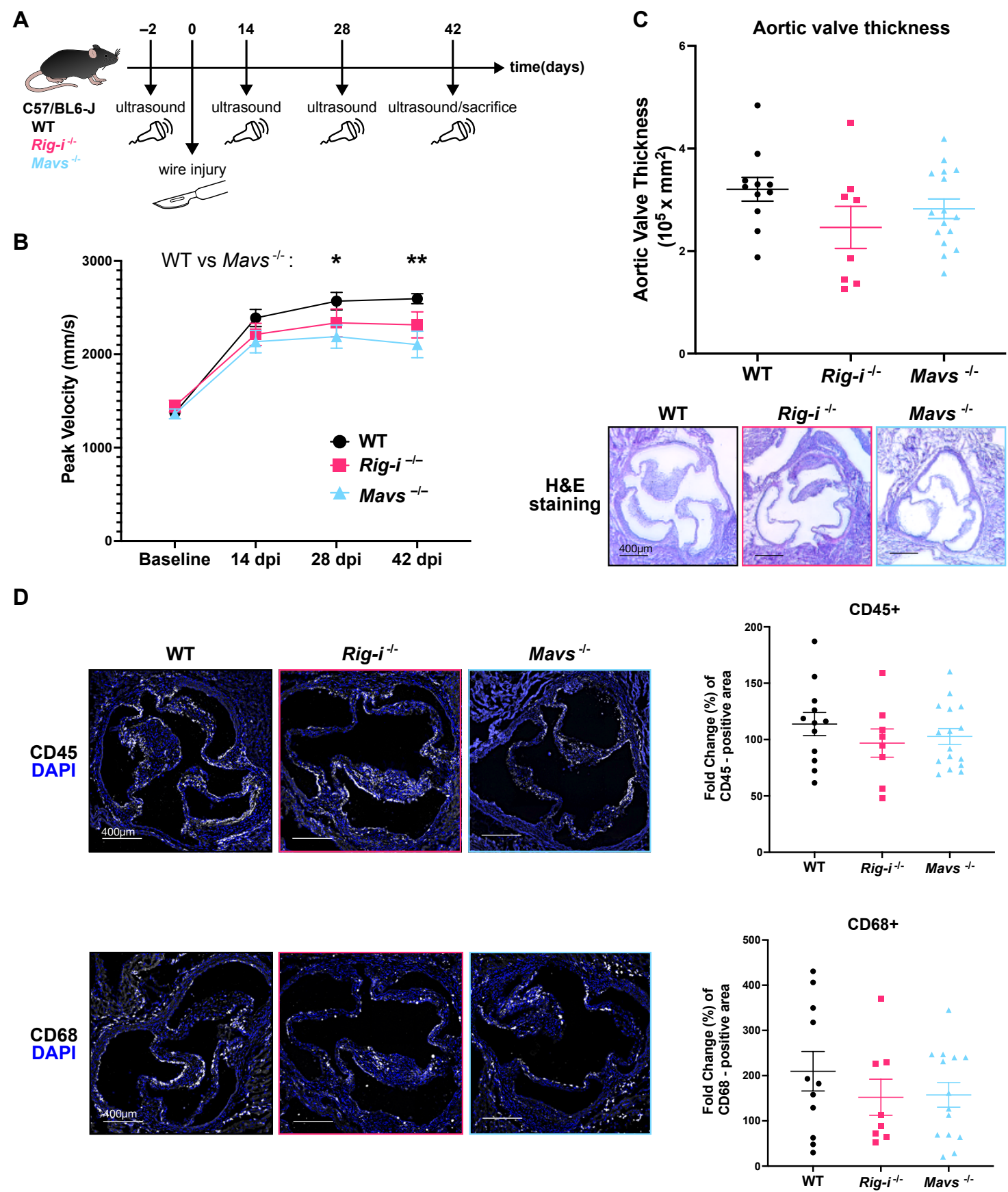

**Supplementary Figure 4: Functional contribution of constitutive RIG-I expression and downstream signaling molecule MAVS to wire-injury-induced aortic valve stenosis.** (A) Schematic overview of the mouse model of wire injury-induced aortic valve stenosis and echocardiographic assessments in WT, *Rig-i*<sup>-/-</sup>, and *Mavs*<sup>-/-</sup> mice on a C57BL/6J background. (B) Trans-aortic valve blood flow peak velocity was determined by echocardiography at baseline and at day 14, day 28 and day 42 post injury (*day post injury*, dpi). Higher peak velocity indicates more severe aortic valve stenosis. (C) Histological analysis of the aortic valve at 42 dpi: representative hematoxylin and eosin (H&E) staining (bottom, scale bar 400 µm) with quantitative analysis (top) of aortic valve thickness (WT n=11, *Mavs*<sup>-/-</sup> n=16, *Rig-i*<sup>-/-</sup> n=8). (D) Representative immunofluorescence staining (left, scale bar 400 µm) of aortic valve sections and quantification (right) of CD45 (leukocytes; WT n=12, *Mavs*<sup>-/-</sup> n=16, *Rig-i*<sup>-/-</sup> n=8) and CD68 (myeloid immune cells; WT n=11, *Mavs*<sup>-/-</sup> n=14, *Rig-i*<sup>-/-</sup> n=8) revealed reduced immune cell infiltration in *Rig-i*<sup>-/-</sup> and *Mavs*<sup>-/-</sup> mice compared to WT mice. Data are shown as individual data points with mean ± SEM. (\* p<0.05, \*\* p<0.01)
