## Supplementary material for "Mechanical stress-induced release of mitochondrial double-stranded RNA activates RIG-I to drive osteogenic differentiation in aortic disease": SupplFig5

**A**

**HAoSMC dsRNA-seq**

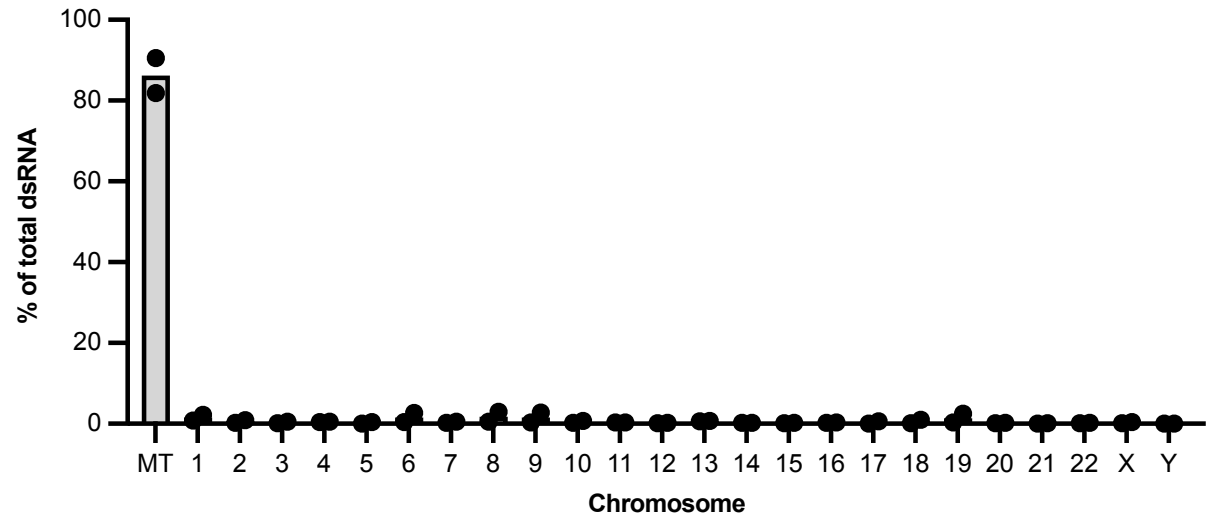

**B**

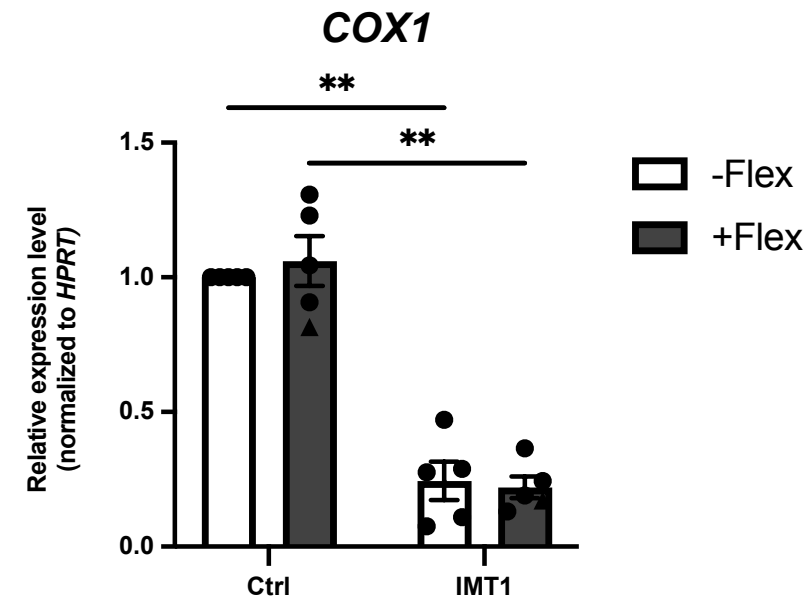

**C**

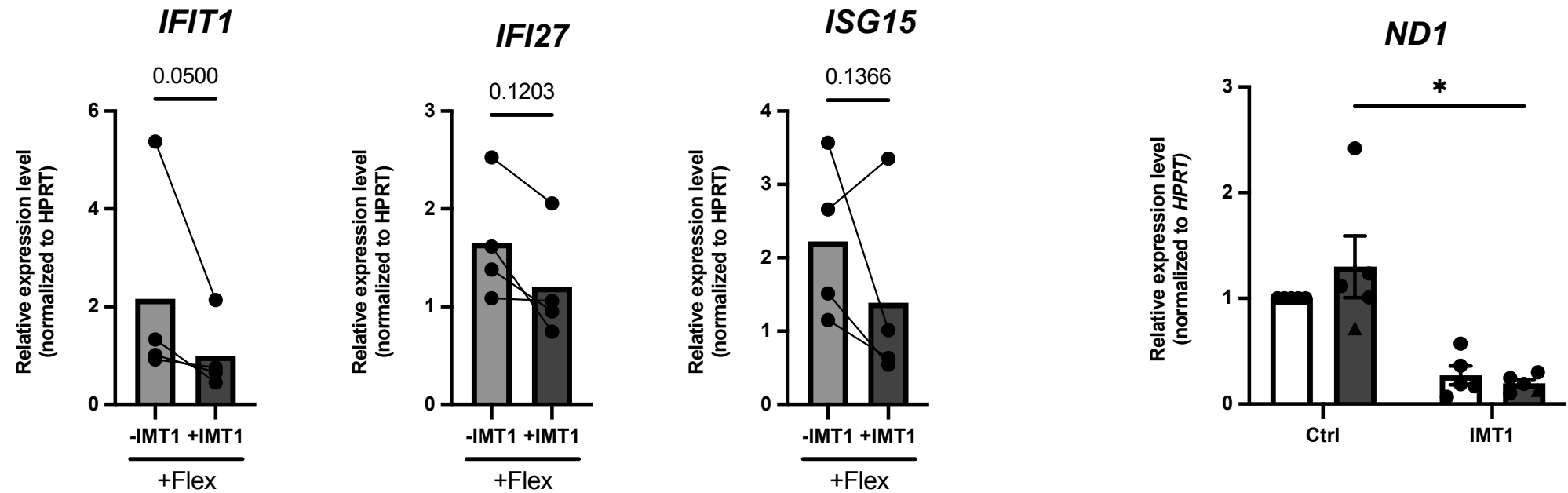

**Supplementary Figure 5: Release of endogenous mitochondrial RNA is responsible for mechanical stress-induced RIG-I activation.** (A) Distribution of total mapped reads after selective sequencing of double-stranded RNA (dsRNA) via SALDOR-Seq in total HAoSMC-derived RNA revealed a predominant mitochondrial origin of endogenous dsRNA species (n=2 donors). (B) HAoSMC were treated with an inhibitor of mitochondrial RNA polymerase (IMT1) 4 days prior to exposure to cyclic mechanical stretch and continued during the exposure to cyclic mechanical stretch (FlexCell Tension System, 0.5 Hz, 15 % elongation, 72 h). Reduced expression of mitochondrial-encoded transcripts (*COX1*, *ND1*; quantified by qRT-PCR) confirmed effective inhibition of mitochondrial RNA transcription upon IMT1 treatment (n=5 donors). (C) HAoSMC treated with IMT1 as in B (mitochondrial RNA depleted) showed reduced expression of *IFIT1* (left), *IFI27* (middle) or *ISG15* (right) upon mechanical stretch (n=4 donors). Data were normalized to untreated control cells. Data are shown as mean ± SEM, each point represents one donor. (\* p<0.05, \*\* p<0.01, \*\*\* p<0.001, \*\*\*\* p<0.0001, ns = not significant; two-way ANOVA with Sidak's multiple comparison test (B) or ratio paired t-test (C)).
